# A Systematically Optimized Miniaturized Mesoscope (SOMM) for large-scale calcium imaging in freely moving mice

**DOI:** 10.1101/2024.02.19.581043

**Authors:** Yuanlong Zhang, Lekang Yuan, Jiamin Wu, Tobias Nöbauer, Rujin Zhang, Guihua Xiao, Mingrui Wang, Hao Xie, Qionghai Dai, Alipasha Vaziri

**Author notes:** Correspondence should be addressed to Q.D. and A.V. These authors contributed equally to this work (Yuanlong Zhang, Lekang Yuan).

## Abstract

Understanding how neuronal dynamics gives rise to ethologically relevant behavior requires recording of neuronal population activity via technologies that are compatible with unconstrained animal behavior. However, realizations of cellular resolution head-mounted microscopes for mice have been based on conventional microscope designs that feature various forms of ad-hoc miniaturization and weight reduction measures necessary for compatibility with the weight-limits for free animal behavior. As a result, they have typically remained limited to a small field of view (FOV) or low resolution, a shallow depth range and often remain susceptible to motion-induced artifacts.

Here, we present a systematically optimized miniaturized mesoscope (SOMM), a widefield, head-mounted fluorescent mesoscope based on a principled optimization approach that allows for mesoscale, cellular resolution imaging of neuroactivity while offering robustness against motion-induced artifacts. This is achieved by co-optimization of a compact diffractive optical element and the associated computational algorithm under form-factor and weight constraints while maximizing the obtainable FOV, depth of field (DOF), and resolution. SOMM enables recordings of neuronal population activity at up to 16 Hz within a FOV of 3.6 × 3.6 mm^2^ in the cortex of freely moving mice while featuring 4-µm resolution, a DOF of 300 µm at a weight of less than 2.5 g. We show SOMM’s performance of recording large-scale neuronal population activity during social interactions, during conditioning-type experiments and by investigating neurovascular coupling using dual-color imaging.

## Introduction

To understand how brain-wide dynamics of highly interconnected neurons in the mammalian brain give rise to ethologically relevant behavior, large-scale neuronal imaging techniques compatible with free behavior and animals engaged in social interactions are required [1]. Recent development of optical imaging techniques in combination with a growing palette of genetically encoded calcium indicators have enabled observing large-scale neuronal population activity in the mammalian brain at the cellular level [2-5]. While these approaches have started to provide a better understanding of how sensory information is represented and transformed across different cortical regions [6, 7], the size and weight constraints associated with traditional desktop microscopes are intrinsically incompatible with ethologically relevant behavioral paradigms. Furthermore, the lack of proprioceptive feedback under head fixation limits the broader applicability, and potentially the validity, of biological findings [8].

On the other hand, recently a number of head-mounted miniaturized microscopes have been developed, allowing access to the most superficial layers of the brain over a sub-millimeter field of view (FOV) [9, 10], and thus enabling discoveries in emotion modulation [11], memory consolidation [12], sleep regulation [13], and ensemble dynamics [14]. To extend the size of the imaged FOV, various technical strategies have been devised including the use of multi-element-lens systems [15-18], low-NA optics [17], and multiplexing [19]. This has come at the cost however, of increased device dimensions, decreased resolution, and increased weight, respectively. Moreover, due to a typically low depth of field (DOF) [10], these image-forming systems are also vulnerable to motion-induced artifacts arising from axial focus drift [20] during free behavior. While components such as electro-tunable lenses and other devices [21] allow for quick adjustment of focus and working distance without mechanical repositioning [16], active compensation for motion-induced axial focus drift during population recordings remains challenging. Thus, to date, these and other technical difficulties have prevented the realization of image-forming devices that offer multi-millimeter FOV, cellular resolution, tolerable weight, and robustness against axial drift in freely behaving rodents.

In this context, a few strategies based on non-image forming optics in combination with image reconstruction and signal extraction algorithms have been demonstrated [22, 23]. While light field microscopy (LFM) [24] represents a special case within this general class of approaches, recently more general types of optical configurations using diffractive optical elements (DOE) have taken the place of conventional lenses, allowing ultra-thin and light-weight form factors [25, 26]. Introducing DOEs in an optical system can relax the demands put on optical design reduce the number of optical components and therefore decrease the weight. These changes are enabled by shifting some of the complexity into the computational realm (via an image retrieval algorithm) which allows for an improvement in overall performance. While initial implementations of this approach have been demonstrated, they either remain limited to small sub-millimeter FOVs offered by gradient-index objectives [27-29], or they offer larger FOV access at the cost of reduced resolution and signal-to-noise ratio (SNR) [30, 31]. Approaches based on microlens arrays as the sole imaging component offer near-cellular resolution across a ∼7 mm FOV, but they are not optimized for freely-moving applications due to their inadequate form factors, weight, and lack of demonstrated practical functionality in freely moving animals [32, 33]. Overall, the tradeoffs between FOV, resolution, and SNR discussed above have limited the practical application of non-image-forming DOE-based devices to large-scale and single-neuron-resolved recordings in freely behaving rodents.

Here, we overcome these limitations for the first time, by principled optimization of hardware and software designs for a non-image-forming, miniaturized, head-mounted mesoscope, thus enabling large-scale, cellular-resolution imaging of neuronal activity in freely behaving mice. Considering the tradeoff between FOV, magnification, resolution, sampling ratio, and form factor, we first determine the manifold of the solution space for an ideal lens and numerically search the maximally obtainable FOV while minimizing the form factor (Fig. 1a). The performance of an ideal lens is experimentally not directly attainable and approximating it would require a prohibitively heavy multi-element optical design. Instead, we took advantage of the flexibility offered by DOEs, developed a joint and iterative design strategy for an optimal DOE and corresponding deconvolution algorithm. We iteratively optimized the performance of the joint system informed by simulations of a detailed and anatomically realistic brain tissue model. This allowed us to identify the DOE phase pattern that permits maintaining the FOV and resolution of a theoretical ideal lens design in practical neuronal population recordings (Fig. 1b). Furthermore, in the optimization process, we required a five-fold increase of the DOE’s depth of field compared to an ideal lens to enhance robustness against axial drift and motion artifacts during free-behavior imaging. Finally, we maximized the illumination uniformity across our mesoscopic FOV by optimizing the distances, orientation angles, and number of illumination sources using a Monte-Carlo method, with the additional aim of enabling two-color imaging without increasing module height (Fig. 1c) as well as optimized the data and power cable to achieve maximum flexibility while retaining high signal transmission efficiency.

**Figure 1.**
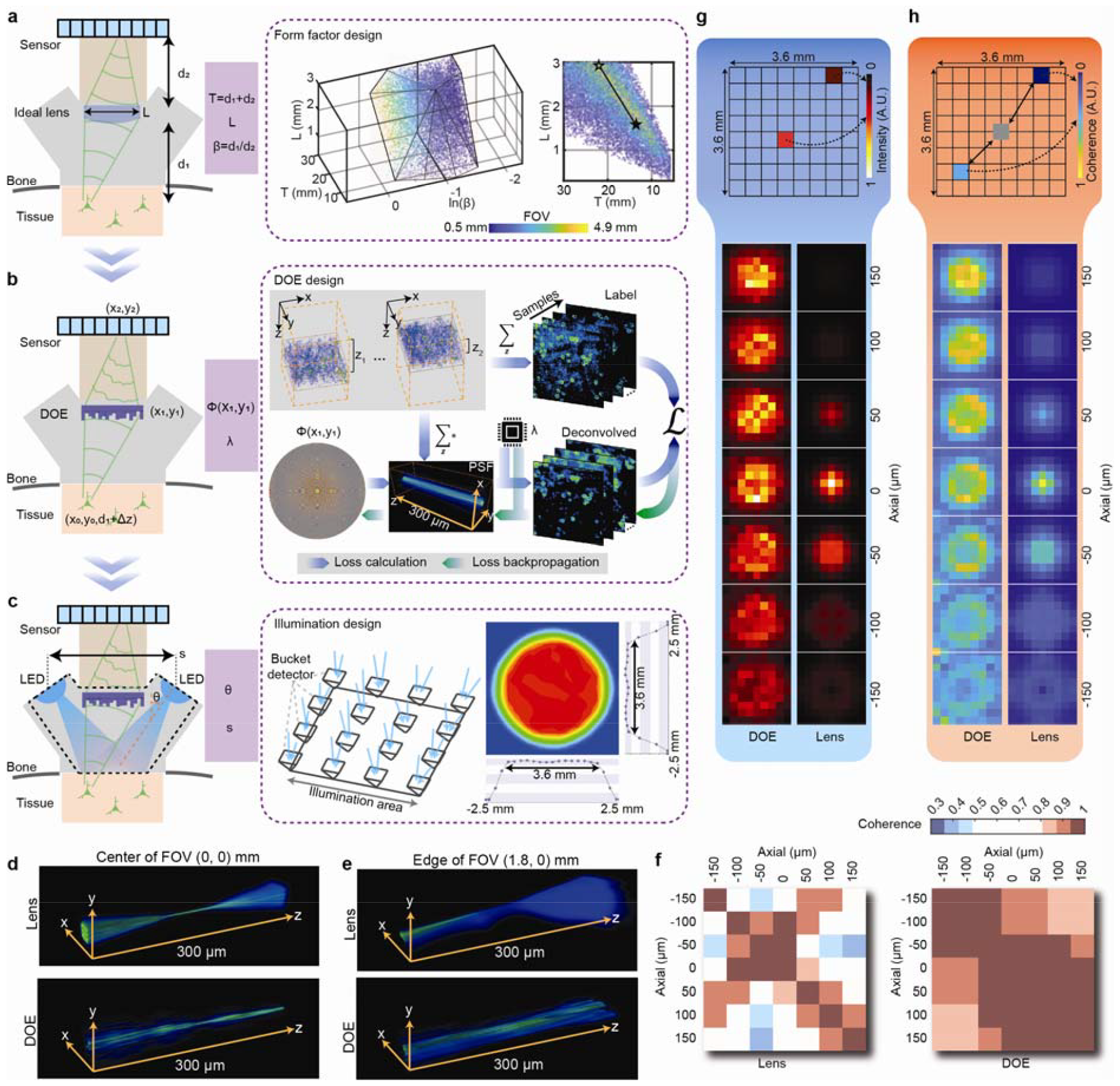
Principle of the systematically optimized miniaturized mesoscope (SOMM). **a**. First optimization round of SOMM: A minimal form factor and maximal field-of-view (FOV) for an ideal imaging system were identified through a joint search of the parameter space while satisfying constraints for feasible angles of light incidence onto pixels, neuron-resolving resolution, and a pixel sampling ratio to fulfill the Nyquist’s limit (Methods). The optimized variables include sample-to-lens distance *d*_1_, lens-to-sensor distance *d*_2_, and imaging aperture *L* To simplify the optimization, the equivalent variables *T = d*_1_ + *d*_2_, β *= d*_1_ / *d*_2_ and were optimized instead. The feasible solutions of *T*, β, and *L* were scattered and formed a manifold where different colors represented different FOVs (from 0.5 mm to 4.9 mm). **b**.Second optimization round of SOMM: The parameter space for a joint solution for diffractive optical elements (DOE) and corresponding deconvolution algorithms that approximated the ideal lens scenario in a. was searched. To achieve this, we fed simulated volumetric neuronal tissue to the DOE-based imaging system via non-paraxial propagation (forward model, blue arrows; Methods), and computed the loss between deconvolved captures and depth-averaged virtual tissue (Methods). The loss was then back-propagated to update both DOE *ϕ*(*x,y*) phase and parameter *λ* of the deconvolution algorithm (backward model, green arrows). The simulated volumetric samples were generated using NAOMi (Methods), and randomly shifted across a 3.6 × 3.6 mm^2^ FOV and 300 μm DOF to resemble large-scale imaging conditions during free behavior with axial drift (Supplementary Fig. 1). **c**. Third optimization round of SOMM: The illumination configurations for uniform illumination in mesoscopic imaging was optimized. Two LEDs that were separated by distance and oriented at angle *θ* were simulated to uniformly emit rays, virtual detectors in the sample plane received those rays and illumination uniformity was evaluated (Methods). After optimization, SOMM achieved uniform illumination across 3.6 mm FOV. **D**. 3D rendering of the point-spread function (PSF) of a plano-convex lens (top) and the optimized DOE (bottom) at the center of the FOV. The plano-convex lens had the same focal length and aperture as the optimized DOE in SOMM. **e**. 3D rendering of PSF of the plano-convex lens (top) and the optimized DOE (bottom) at the edge of the FOV (x = 1.8, y = 0 mm from the center). **f**. Cross-coherence matrix of PSFs across different depths for the plano-convex lens (left) and the optimized DOE (right). The cross-coherence was calculated between PSFs from each depth, with specific depths labeled on the side. The details of the calculation can be found in Methods. All panels share the same color bar displayed at the top. **g**. Peak intensities of PSFs of the optimized DOE (left) and the plano-convex lens (right) across 3.6 × 3.6 mm^2^ FOV. As illustrated at the top, the whole FOV was tiled into a 9 × 9 grid for investigating peak intensities of PSFs for different positions in the FOV. All panels share the same color bar displayed at the top. **h**. Cross-coherence of PSFs by the optimized DOE (left) and the plano-convex (right) across 3.6 × 3.6 mm^2^ FOV. As illustrated at the top, the whole FOV was tiled into a 9 × 9 grid for investigating peak intensities of PSFs for different positions in the FOV. All panels share the same color bar displayed at the top. The cross-coherence was calculated between the PSF at each position and the PSF at the center, as visualized at the top. Details of the calculation can be found in Methods.

Collective implementation of these optimization measures allowed us to realize a systematically optimized miniaturized mesoscope (SOMM) capable of mesoscale imaging of neuroactivity across a 3.6 × 3.6 mm FOV at 4 µm lateral resolution and over a 300 µm DOF at a device weight of only 2.5 grams. Thus, SOMM enables largescale cellular-resolution imaging and recording of neuronal population activity at 16 Hz volume rate in freely behaving mice (Supplementary Table 1). We verified SOMM’s performance by recording cortex-wide neuronal activity in freely behaving mice under various conditions. This includded during free exploration of an arena, engagement in uninstructed social interactions, and fear-conditioning experiments using visual cues, at single-neuron resolution and across tens of cortical regions spanning the visual, retrosplenial, and somatosensory areas. We also demonstrated the potential of SOMM to reveal interactions between neuronal population activity and dynamic properties of vascular networks via dual-color imaging.

## Results

### Comprehensive optimization for SOMM

To overcome the current technical limitations in realizing a large-scale and light-weight cellular resolution calcium imaging system compatible with imaging in freely having mice, we developed and implemented a systematic and comprehensive optimization scheme for our hardware and software. This process consisted of three key steps and was aimed at identifying an optimal tradeoff between resolution, FOV, SNR, robustness against axial drift, form factor, and weight.

As the first step we used an ideal lens approximation and sought to identify the solution space that minimized the form factor while maximizing the FOV. Mathematically, the optimization was carried out over the lens-sample distance *d*_1_, sensor-lens distance *d*_2_, lens diameter *L*, constrained by the feasible angle of ray incidence onto the pixels [34], neuron-resolving resolution, and the requirement that the size of the magnified resolvable spot size should extend over more than two camera pixels, thus satisfying the Nyquist limit (Methods; Supplementary Note 1). The solution space of possible combinations of these parameters is shown in Fig. 1a. We found that the largest FOV satisfying these constraints was ∼4.9 mm, resulting in a total axial form factor *T* = *d*_*1*_+*d*_*2*_ of ∼23 mm, achieved by *d*_*1*_*≈d*_*2*_and ≈2.9 mm. However, examining the tradeoff between *T* and FOV we found that the axial footprint can be reduced by more than 40% while sacrificing only less than 8% in FOV (black line and star in Fig. 1a) by reducing the lens diameter *L*. Thus, we settled on overall values for our design parameters of T=*d*_*1*_+*d*_*2*_ ≈ 14mm, *L* ≈ 1.5 m with a target FOV of ∼4.5 mm. These parameters lead to a ∼40% smaller axial footprint and a FOV up to ∼50 times larger for our system compared to the popular Miniscope v3 system without the sensor [10].

The above examination of basic tradeoffs between design parameters assumes an ideal lens, that is, it does not consider the aberrations that would be expected at such a large FOV. Thus, in a second step, we substituted the ideal lens with a combination of a DOE and a deconvolution algorithm. Specifically, we iteratively and jointly optimized the phase profile (*ϕ*) of the DOE and the parameters (*λ*) of the deconvolution algorithm to approximate the ideal lens design while maximizing excitation efficiency [31] and uniformity of optical performance across the FOV (Fig. 1b). Furthermore, the flexibility provided by this approach allowed us to achieve the above specifications while extending the axial point spread function (PSF) of our DOE, thereby mitigating axial motion artifacts during vigorous animal movement; a common issue with conventional miniaturized microscopes. By optimizing the DOE to an axial PSF of about five times longer than the corresponding diffraction-limited PSF (60 µm at 1/e full width) across the target FOV, individual neurons could still be resolved in the presence of axial motion events and their corresponding activities could be temporally demixed for intermediate density of neuronal labeling. To achieve the above iterative optimization between the parameters of the DOE and the deconvolution algorithm, we employed an optically informed convolutional neuronal network (CNN), conceptually similar to a recently described technique [35, 36]. We tailored the algorithm based on an angular spectrum method instead of the original Fresnel propagations to maximize the fidelity in simulating the system’s PSF (Methods). We generated highly realistic cortical volumes of 200 × 200 × 60 µm using the method of Neural Anatomy and Optical Microscopy (NAOMi) [37], and applied random axial shifts of up to ±150 µm to intentionally introduce defocusing during training and thus encourage defocus-robust detection (Supplementary Fig. 1b). Equipped with both the optical kernels and the realistic sample models, we then produced a forward model (blue arrows in Fig. 1b) for our imaging system by convolving our realistic cortical volume with candidate PSFs and projecting them along the axial (*z*) direction to simulate a sensor image. Finally, we employed a deconvolution algorithm with learnable parameters (*λ*) to restore the original volume (Methods). The L2 loss between deconvolved images and the axial summation of the brain tissue images (Supplementary Fig. 1c), the PSF concentration loss [38], and vignetting loss (Methods) were back-propagated (using automatic differentiation of the CNN) to update both the DOE phase profile *ϕ* and the deconvolution parameters *λ* (green arrows in Fig. 1b, Supplementary Fig. 2). Iteration of the above procedures led to an optimized DOE that facilitated uniformly high-contrast and localized PSFs across a 3.6 × 3.6 mm^2^ FOV (Fig. 1d, e, Supplementary Fig. 3), and also resulted in corresponding optimal parameters of the deconvolution algorithm. Since the above process requires optimization of a corresponding deconvolution parameter for each lateral FOV position, the desired uniform imaging performance across the FOV cannot be achieved using traditional shift-invariant deconvolution algorithms (Supplementary Fig. 4). We therefore developed a new shift-variant deconvolution (SV-Deconv) algorithm which utilizes different PSFs and deconvolution parameters for different lateral patches of the FOV, resulting in patches with sharp features for all patch locations, and then concatenates these patches to form globally sharp images (Methods). We demonstrated that SV-Deconv facilitates uniform imaging performance across the entire FOV, outperforming traditional shift-invariant deconvolution with regard to various image quality metrics, including the well-known PSNR, SSIM, and RMSE metrics (Supplementary Fig. 4).

**Figure 2.**
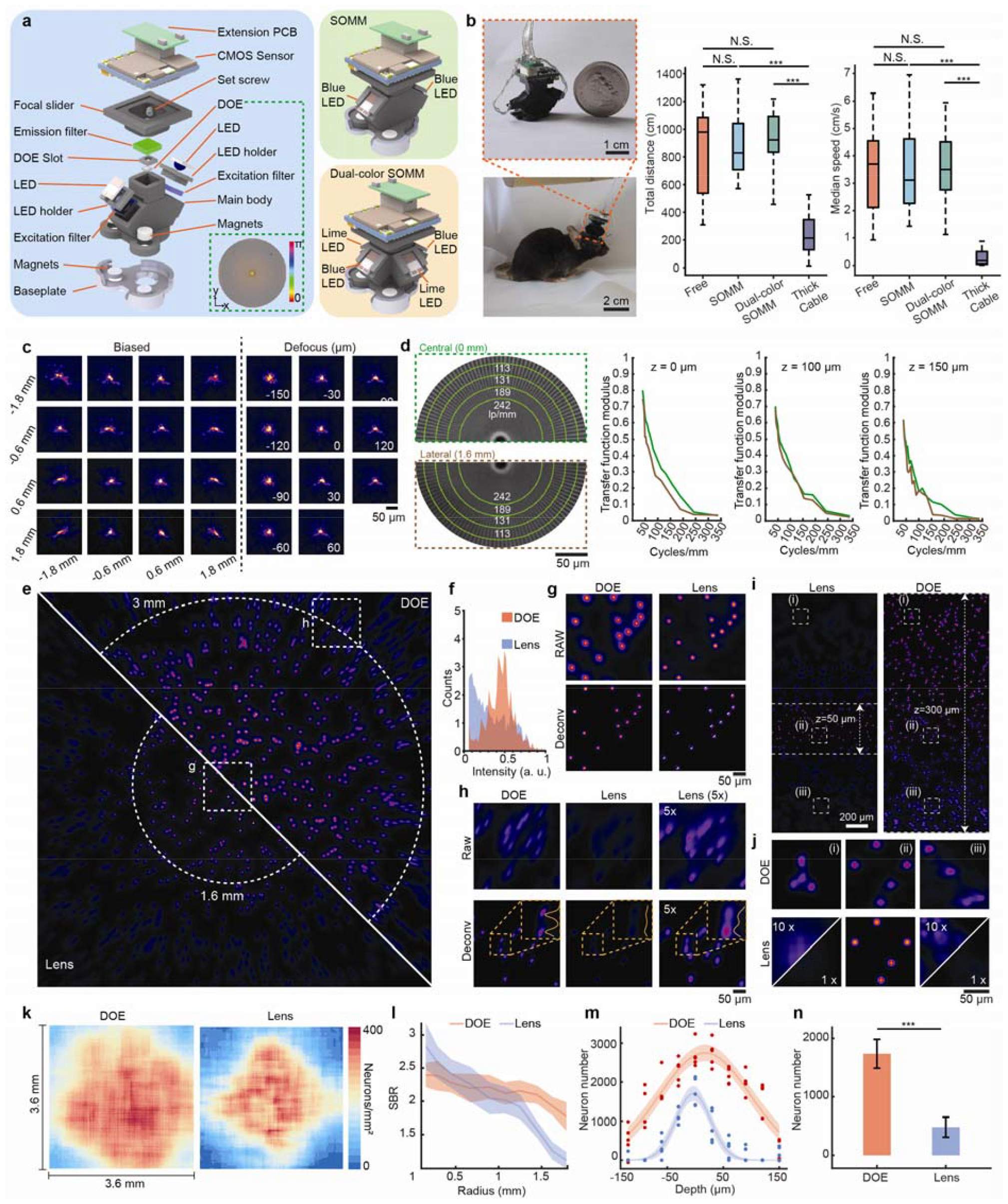
SOMM performance validation. **a**.3D rendering of SOMM assembly. Left, explosion diagram of SOMM. The DOE phase profile wrapped into [0,*π*] is illustrated in the bottom right corner. CMOS, complementary metal-oxide semiconductor. PCB, printed circuit board. LED, light-emitting diode. DOE, diffractive optical element. Right, 3D rendering of SOMM (top) and dual-color SOMM (bottom). **b**. Left, a still image of a mouse carrying SOMM and engaging in natural behavior. Top: Zoom-in panel: side-by-side comparison of a full-assembled SOMM and a quarter dollar coin. Right, box plot showing total distance and median speed when the mice traveled in a home cage (20 cm wide) in conditions without the head-mounted mesoscope (free), with SOMM connected by an optimized cable (SOMM), with dual-color SOMM connected by an optimized cable (dual-color SOMM), and with SOMM connected by a non-optimized cable (thick cable). Each condition was tested in a 3-minute session. Center line, median; limits, 75% and 25%; whiskers, maximum and minimum. Conditions are compared using one-way ANOVA test where N.S. > 0.1 and ^***^p < 0.001. **c**. Calibrated PSF stacks of SOMM measured across 3.6 × 3.6 mm^2^ FOV (left, at z = 0 µm) and 300 μm DOF (right, at positions with x = 0 mm and y = 0 mm). **d**. Characterization of the modulation transfer function (MTF) of SOMM. Left, a segment of deconvolved Siemens star chart captured with SOMM. In this segment, line density varies from 113 to 242 line pairs per mm (8.8 to 4.1 µm spacing). Top: chart at the center of FOV, bottom: chart measured at the edge of FOV (1.6 mm from the center). Right, measured MTF at the center of FOV (green) and at the edge of FOV (1.6 mm from the center, brown) at z = 0 µm, z = 100 µm, and z = 150 µm, respectively. **e**. Images of fluorescent beads (diameter 6 μm) taken with SOMM (labelled “DOE”, top right) and a replica with a plano-convex lens (“Lens”, bottom left; Methods). The dashed white circle encloses the area with a relative illumination intensity greater than 0.4. **f**. Intensity histograms of fluorescent beads detected by SOMM (“DOE”, red) and the replica with a plano-convex lens (“Lens”, blue). **g**.Zooms into the white dashed box that is close to the optical axis in e by SOMM (“DOE”, left column) and the replica with a plano-convex lens (“Lens”, right column). Top row: raw captured images. Bottom row: images deconvolved by SV-Deconv algorithm. **h**. Zooms into the white dashed box that is close to the edge of FOV in e by SOMM (“DOE”, left column), the replica with a plano-convex lens (“Lens”, middle column), and the image from the replica shown at 5× brightness (right column). Top row: raw images. Bottom row: images deconvolved by SV-Deconv algorithm. In the bottom row, intensity profiles across two fluorescent beads are plotted in yellow. **i**. Left, tilted fluorescent beads sample (diameter 6 μm) imaged by the replica with a plano-convex lens (“Lens”, left; Methods) and SOMM (“DOE”, right). The slide was tilted by 21.6 degrees such that the axial positions of fluorescent beads were mapped to their different lateral positions (Supplementary Fig. 14). The dashed white line encloses the area with relative illumination intensity greater than 0.4. **j**. Zooms into white dashed boxes in i for SOMM (“DOE”, the first row) and the replica with a plano-convex lens (“Lens”, the second row). Images were deconvolved by the SV-Deconv algorithm. The defocused fluorescent beads in the lens-based replica images are shown at 10× intensity for clarity. **k**. Detectable neuron density in tissue by SOMM (“DOE”, left) and the replica with a plano-convex lens (“Lens”, right) across 3.6 × 3.6 mm^2^ FOVs. The tissue was simulated using NAOMi (Methods). Results were averaged over n = 5 simulation runs. **l**.Signal-to-background ratio (SBR, Methods) of extracted neurons from SOMM data (“DOE”, red) and the replica with a plano-convex lens (“Lens”, blue), as a function of lateral distance to the optical axis. Shaded area: SD, solid line: mean. **m**. Detectable neuron number across 300 μm axial defocus range for SOMM (“DOE”, red) and the replica with a plano-convex lens (“Lens”, blue). Data points: n = 5 samples for each axial depth. Solid line: exponential fit, shaded area: 95% confidence interval of fit. **n**. Overall number of detected neurons across 3.6 × 3.6 mm^2^ for SOMM (“DOE”, red): 1736 ± 123 (mean ± SE), compared to 481 ± 85 (mean ± SE) for the replica with a plano-convex lens (“Lens”, blue), n = 5 samples. Height: mean. Error bars: 95% confidence interval of the mean. ^***^p < 0.001, two-sided Wilcoxon signed-rank test.s Scale bar, 1 cm (top) and 2 cm (bottom) in **a**, 50 µm in **c, d, g, h**, and **j**, and 200 µm in **i**.

**Figure 3.**
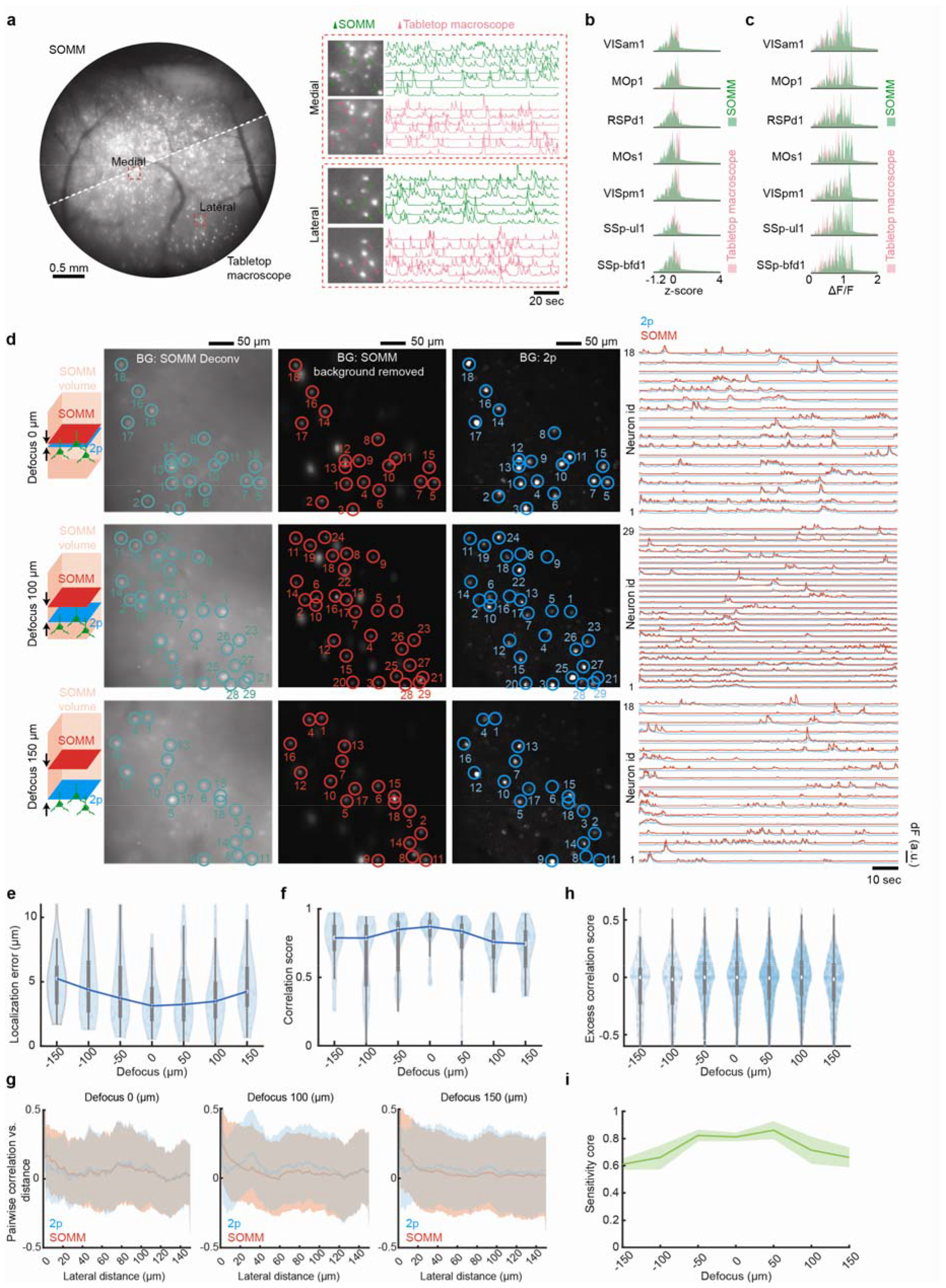
Verification of SOMM against tabletop one-photon widefield system and two-photon microscope. **a**. Spatial and temporal comparison between SOMM and a tabletop widefield imaging system. Left, Standard deviation projection (STD) of a deconvolved SOMM video (top left) and STD of a tabletop widefield system video (bottom right). Both were captured at the same speed (16 Hz) and for 5 minutes. Right, zoom-in areas in medial (top, 0.6 mm from the FOV center) and lateral (bottom, 1.3 mm from the FOV center) regions of the FOV, with corresponding neuronal activities plotted on the right side. Green markers and traces: SOMM. Magenta markers and traces: tabletop one-photon widefield system. **b**. Comparisons of z-score values of neuron activities for SOMM and tabletop one-photon widefield system. Histogram of z-scores of neuron activities for a 5-minute video captured with SOMM (green) and tabletop one-photon widefield system (magenta) across multiple brain regions. **c**. Comparisons of ΔF/F values of neuron populations for SOMM and tabletop one-photon widefield system for 5-minute video captured with SOMM (green) and the tabletop one-photon widefield system(magenta) across multiple brain regions. **d**. Spatial and temporal evaluation of SOMM with 2p functional ground truth. Temporally interleaved 2p functional ground truth was generated by interleaving SOMM frames and planar 2p microscope frames (Supplementary Fig. 18a, Methods). First column: schematic plot showing the relative positions of 2p (blue) and SOMM (red) focal planes within the SOMM detection volume (pink). Second to forth columns: STD of deconvolved SOMM video, STD of background removed SOMM video through DeepWonder [39], and STD of 2p video, for 3 different SOMM defocus configurations (top: 0 μm; middle: 100 μm; bottom: 150 μm; see details on defocus configuration in Supplementary Fig. 18b). SOMM-extracted neuronal positions are shown as circles overlaid on the deconvolved (second column, green) and background-subtracted videos (third column, red). Paired 2p neuronal positions were analyzed by CaImAn followed by human annotation, and were overlaid in the 2p videos (forth column, blue circles). Fifth column: neuronal activity traces from SOMM (analyzed by the proposed pipeline, red) and 2p (analyzed by CaImAn, blue) corresponding to circles in left panels, as used for performance quantifications. **e**. Distributions of lateral neuron localization errors between SOMM-extracted neuron positions and experimental functional temporally interleaved ground truth. White circle: median. Thick grey vertical line: Interquartile range. Thin vertical lines: upper and lower proximal values. Transparent blue disks: data points. transparent violin-shaped area: kernel density estimate of data distribution. n = 439 neuron pairs from 4 recordings. **f**. Distributions of temporal correlations between experimental “temporally interleaved” ground truth activity traces and matched SOMM traces versus different 2p-SOMM focal plane distances. Violin plot elements as in e. n = 439 neuron pairs from 4 recordings. **g**. Mean pairwise correlation between all pairs of traces in “temporally interleaved” experimental functional ground truth (blue line) and mean pairwise correlation between corresponding pairs of SOMM-extracted traces (red line) as a function of lateral distance between the neurons in the pairs. Three 2p-SOMM focal plane distances (left: 0 μm; middle: 100 μm; right: 150 μm) are shown. No significant change in pairwise correlation, i.e., in the difference between SOMM-extracted correlation and ground truth correlation, was observable across all pair distances (n = 7943 neuron pairs from 4 recordings). Shaded areas: mean ± SD. **h**. Distributions of excess correlation between pairs of neuronal traces in experimental “temporally interleaved” ground truth and corresponding pairs of SOMM-extracted traces, as a function of 2p-SOMM focal plane distances. For all distances, the modulus of the mean excess correlation is below 0.07, indicating robust crosstalk rejection by the SOMM processing pipeline. Violin plot elements as in e. n = 7943 neuron pairs from 4 recordings. **i**. Neuron detection score sensitivity achieved by SOMM on experimental temporally interleaved functional verification dataset as a function of depth. Shaded areas: mean ± SD; data from n = 4 recordings. Scale bar, 0.5 mm and 20 seconds in a, 50 μm and 10 seconds in **d**.

**Figure 4.**
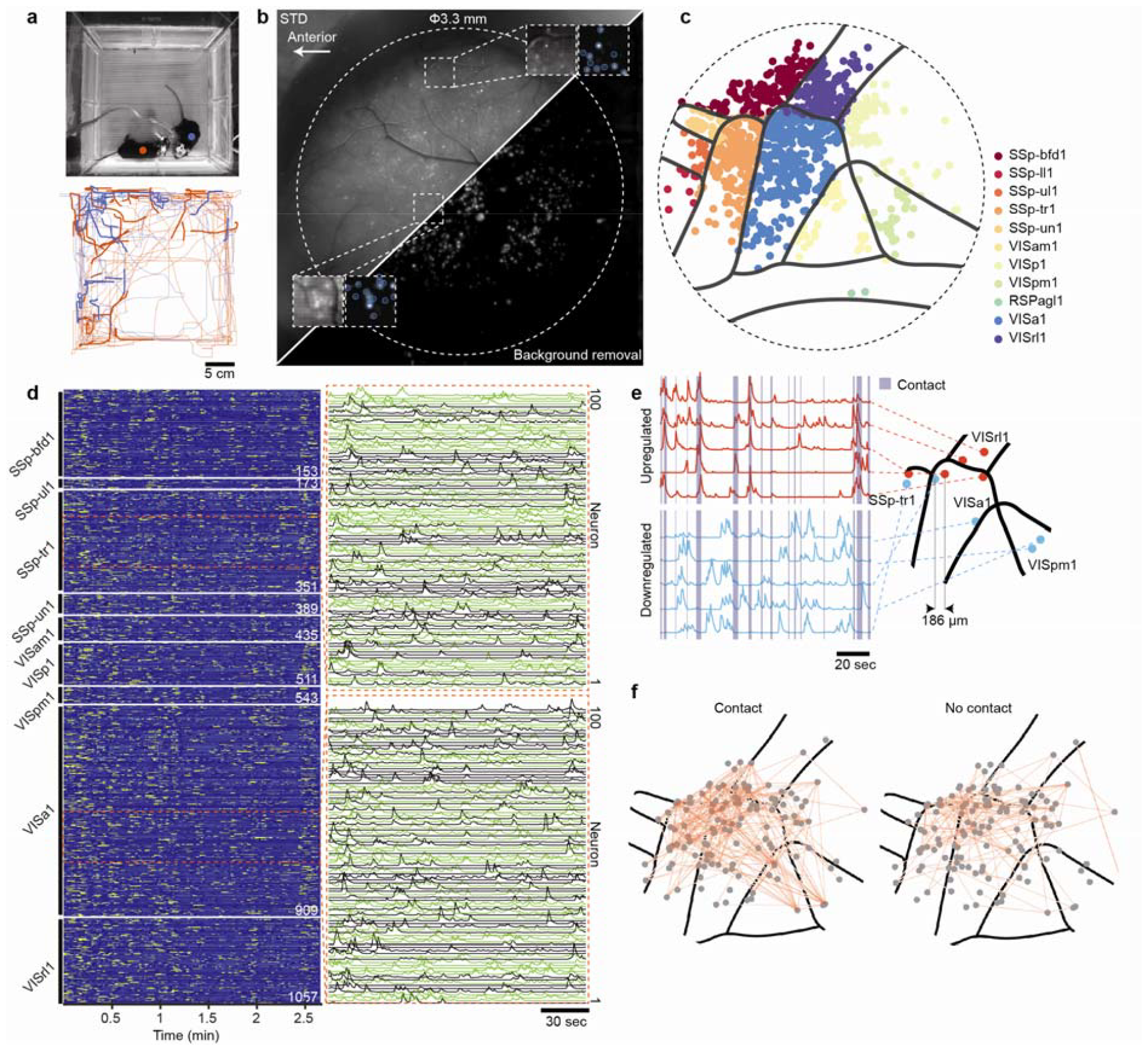
SOMM detects cortex-wide neuronal activities in freely-behaving mice during social interaction. **a**. Top, a photograph of a mouse with SOMM (red dot) interacting with a companion mouse (blue dot). Bottom, traces of instantaneous locations of the two mice during the trial. The segments of thicker and darker blue and red indicated the contact, where positions of the two mice were within 5 cm of each other. **b**. Maximum intensity projection (MIP) of deconvolved SOMM movie (top left) and MIP of background removed movie by the processing pipeline (bottom right, Methods). White dashed boxes marked zoom-in one central and one marginal area of FOV in the cortex, with neuron segmentation (blue circles) overlaid. The direction towards the anterior side was labeled by a white arrow. The white dashed circle marked effective FOV which was limited by the size of the cranial window. **c**. Segmented neurons in **b** overlaid with Allen CCF atlas [49] with different colors representing different cortical regions. **d**. Temporal activity rendering of 1057 neurons inferred by the processing pipeline in a 2.7-minute recording. Two zoom-in panels showed example traces (each with 100 neurons). **e**. Tuning analysis in a single-cell level. Left, temporal activities of neurons that were upregulated (red) and downregulated (blue) by social interactions. Deep blue shadows represented the periods when two animals touched each other. Right, spatial distributions of those neurons aligned in the CCF atlas. **f**. Functional connection analysis in a single-cell level. Network nodes were neurons that had strong connections (Pearson correlation higher than 0.8) with other neurons. Thickness of network edges represented Pearson correlation strength. Two social interaction states (contact and no contact) were plotted. SSp-bfd1, Primary somatosensory area barrel field layer 1; SSp-ll1, Primary somatosensory area lower limb layer 1; SSp-ul1, Primary somatosensory area upper limb layer 1; SSp-tr1, Primary somatosensory area trunk layer 1; SSp-un1, Primary somatosensory area unassigned layer 1; VISam1, Anteromedial visual area layer 1; VISp1, Primary visual area layer 1; VISpm1, posteromedial visual area layer 1; RSPagl1, Retrosplenial area lateral agranular part layer 1; VISa1, Anterior area layer 1; VISrl1, Rostrolateral area layer 1. Scale bar, 5 cm in **a**, 30 seconds in **d**, and 20 seconds in **e**.

Our optimized DOE holds several key advantages compared to a conventional spherical lens. First, the PSFs generated by our optimized DOE are axially elongated with an invariant profile across the 300-µm depth range (Fig. 1d-e), thus eliminating the need for 3D scanning while accessing axially distributed neurons. Second, the cross-coherence of our PSF between the center of the FOV and a position of 1.5 mm from the center was more than 20 times greater than a conventional lens (at axial position z = 0 µm), while the peak intensity was 10 times higher (Fig. 1g, h), representing significantly improved lateral invariance of our PSFs and reduced vignetting. Third, both the cross-coherence and the peak intensity across different axial positions were considerably higher (Fig. 1f-h), thanks to the extended defocus robustness and detection ability. Overall, we confirmed that our system exhibited a lateral resolution of 4 µm after deconvolution across the 3.6 × 3.6 mm^2^ FOV over ∼300 μm axially (Supplementary Fig. 5). We note that to achieve a similar lateral resolution over such a large FOV, a spherical lens system with ∼130-times more glass volume would be required (Supplementary Fig. 6). Such an optical system would be incompatible with the weight requirements of an instrument for free-behavior neuroimaging and would still lack the axial robustness as achieved by the DOE (Supplementary Fig. 7). Thus, the unique feature of our optimized DOE is that it allows us to obtain a large FOV, single-neuron resolution, robustness against axial drift, and compactness within a single design and realization.

**Figure 5.**
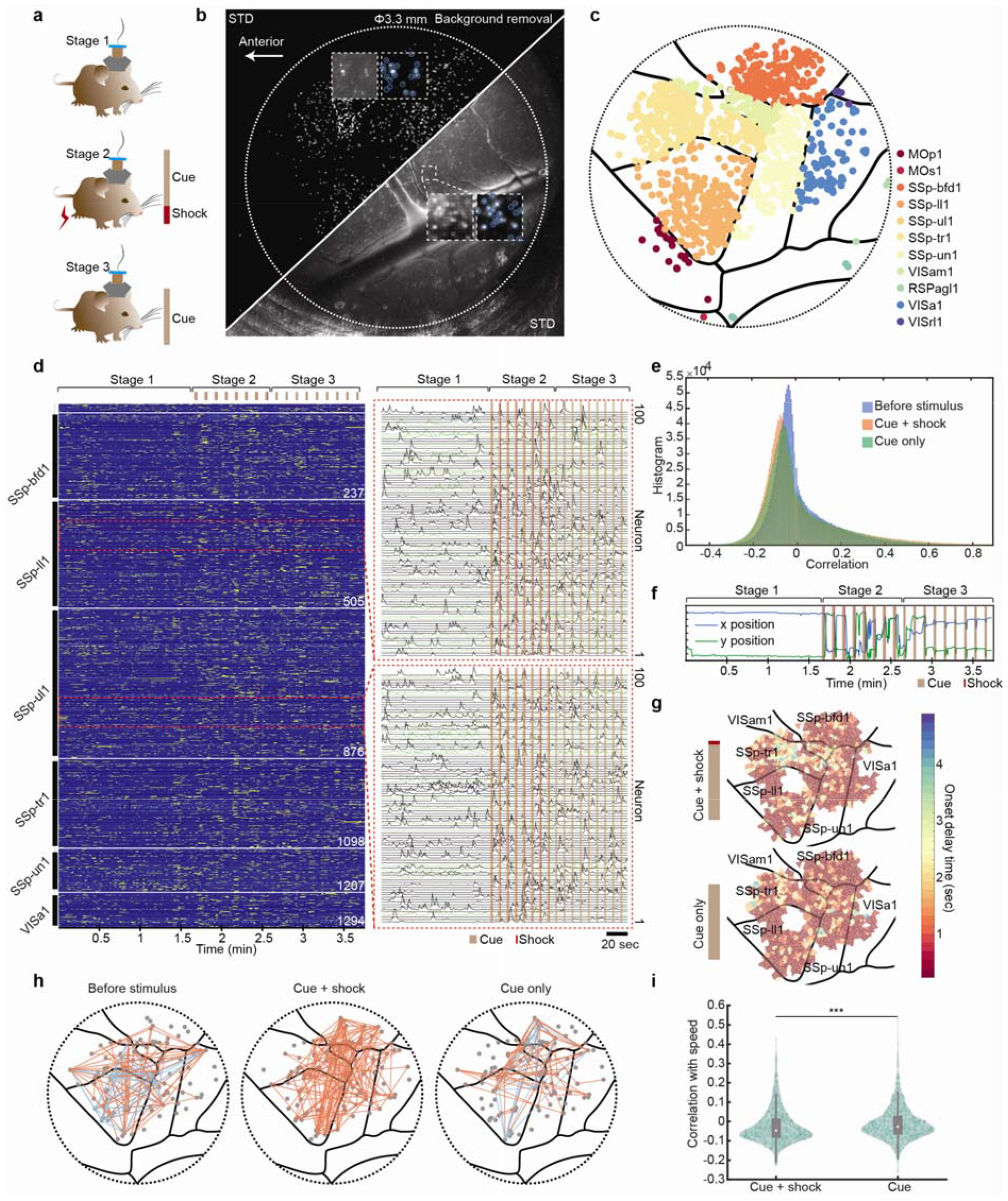
SOMM reveals electrical-stimulus-evoked activities in freely behaving mice across tens of cortical regions. **a**. Conditioning test assay for freely behaving animals. Firstly, animals were freely behaving in an open field (top, 20 cm side). Then, animals received both a visual cue (brown) from a LED and then an electrical shock (red) from the ground (middle). Lastly, animals only received a visual cue from the LED. **b**. Maximum intensity projection (MIP) of deconvolved SOMM movie (top left) and MIP of background removed movie by the processing pipeline (bottom right, Methods). White dashed boxes marked zoom-in areas in central and marginal positions of FOV in the cortex, with neuron segments overlaid. The white dashed circle marked effective FOV which was limited by the size of the cranial window. **c**. Segmented neurons in b overlaid with Allen CCF atlas with different colors representing different cortical regions. **d**. Temporal activity rendering of 1314 inferred neurons in a 3.8-minute recording. Stimuli of cue (brown) and shock (red) were marked in the top and aligned with the neuronal activities. Two zoom-in panels showed example traces (each with 100 neurons) with the stimulus of cue (brown) and shock (red) overlaid as shadows. **e**. Histograms of mutual correlations between recorded neurons during the assay. Different colors represented different periods, where blue was for freely moving before stimuli, orange was for receiving both cues and shocks, and green was for receiving only cues. **f**. Animal instantaneous locations during the assay. The positions consisted of x (blue) and y (green) dimensions and were plotted with stimuli of cues (brown) and shocks (red) overlaid as shadows. **g**. Distribution of the onset delays of cue-evoked activity peaks during the assay. The onset delay time for each of the neurons was counted as the time between the visual stimulus and the arrival of the first peak activity. Conditions with both cues and shocks (top) and only cues (bottom) were both plotted across cortical regions. **h**. Functional connections of the inferred neuronal population during the assay. The functional connections translated during freely moving before stimulus (left), receiving both cues and shocks (middle), and only receiving cues (right). Network nodes were neurons that had strong connections (absolute of Pearson correlation higher than 0.8) with other neurons. Orange for positive correlations and blue for negative correlations. **i**. Violin plots of correlations between neuronal activities and animal moving speed during the assay. Correlations during periods of receiving both cues and shocks (left) and only receiving cues (right) were both plotted. ^***^P<0.001, two-sided Wilcoxon signed-rank test across n = 1314 neurons. White circle: median. Thick grey vertical line: Interquartile range. Thin vertical lines: upper and lower proximal values. Transparent green disks: data points. Transparent violin-shaped area: kernel density estimate of data distribution. MOp1, Primary motor area Layer 1; MOs1, Secondary motor area layer 1; SSp-bfd1, Primary somatosensory area barrel field layer 1; SSp-ll1, Primary somatosensory area lower limb layer 1; SSp-ul1, Primary somatosensory area upper limb layer 1; SSp-tr1, Primary somatosensory area trunk layer 1; SSp-un1, Primary somatosensory area unassigned layer 1; VISam1, Anteromedial visual area layer 1; RSPagl1, Retrosplenial area lateral agranular part layer 1; VISa1, Anterior area layer 1; VISrl1, Rostrolateral area layer 1. Scale bar is 20 seconds in **d**.

**Figure 6.**
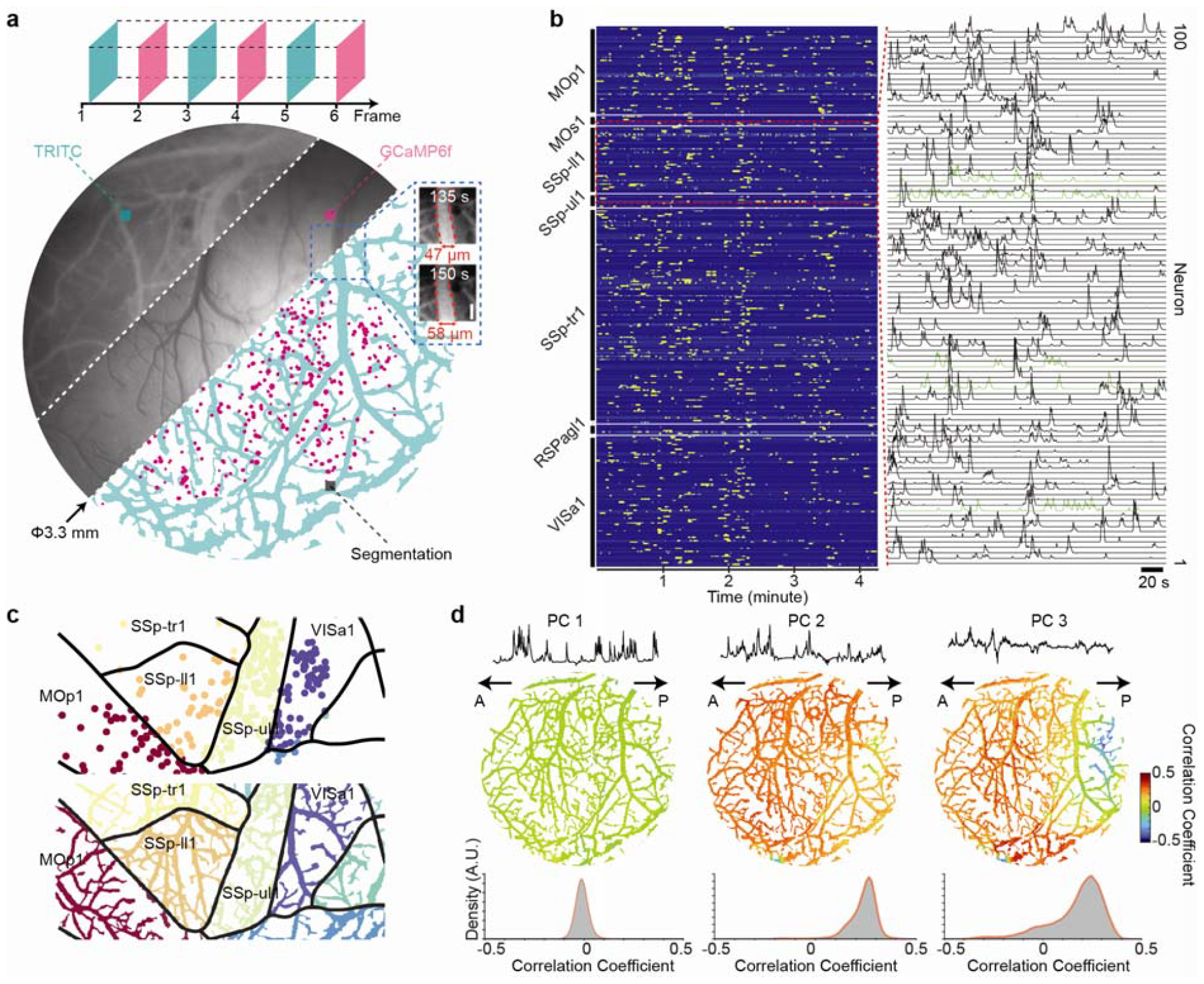
Dual-color SOMM uncovers neurovascular coupling at mesoscopic scale. **a**. Interleaved dual-color recording of neuronal activity (GCaMP6f) and vasculature dynamics (TRITC) using SOMM (Methods). Two groups of LEDs with different colors were switched on alternatingly and in sync with the sensor to separately capture fluorescence in the two different color channels (top). Large circular image is a combination of raw frames of TRITC labeled vessels (top left segment), raw frames of GCaMP6f labeled neurons (middle), and their respective segmentation (bottom right; blood vessels in green and neurons in magenta). Zoom-in panel shows the dilation of a vessel at two time points with vessel diameter labeled (right). **b**. Left: Heatmap of temporal activity of 342 inferred neurons in a 4.3-minute recording. Right: Zoom into the region marked with dashed red lines in the heatmap, showing activity of 100 neurons as stacked line plots. **c**. Segmented neurons (top) and vessels (bottom) overlaid with Allen CCF atlas, with colors indicating different cortical regions. The shown area is a cropped region of the full FOV, for clarity. **d**. Correlation between principal components of neuronal population activity and the vascular dynamics. Top row, three principal components (PCs) of 342 inferred neuronal activity signals. Middle row, heatmaps of correlation coefficients between PCs and vascular dynamics reported by changes of fluorescence due to modulation of blood flow. Bottom row, histograms of correlation coefficients between PCs and vascular dynamics. Anterior (A) and posterior (P) directions are labeled with black arrows. MOp1, Primary motor area Layer 1; MOs1, Secondary motor area layer 1; SSp-ll1, Primary somatosensory area lower limb layer 1; SSp-ul1, Primary somatosensory area upper limb layer 1; SSp-tr1, Primary somatosensory area trunk layer 1; RSPagl1, Retrosplenial area lateral agranular part layer 1; VISa1, Anterior area layer 1. Scale bars: **a**, 50 µm. **b**, 20 seconds.

Finally, we systematically optimized the illumination configuration of our head-mounted mesoscope for maximizing the illumination flatness and minimizing imaging aberrations over the FOV (Fig. 1c). We found that a co-axial illumination configuration with a 45-degree dichroic mirror as used in most conventional miniaturized microscopes would introduce significant wavefront errors in a single-lens imaging system such as SOMM (Supplementary Fig. 8). As an alternative, we designed and implemented a side illumination strategy that avoids aberrations in the imaging path. However, this approach required careful design to suppress illumination non-uniformities across our large FOV. To identify the optimal illumination strategy while minimizing any increase in device form factor, we searched for the optimal choice of the illumination angles (*θ*) for a fixed LED separation distance (*s*) and subsequently searched for the optimal distance s. This was done by simulating randomly oriented illumination rays emanating from LEDs and identifying the parameter for which the ray density across the target FOV was maximized over an array of virtual bucket detectors (Fig. 1c, Methods). The optimal design solution generated using this approach resulted in over 85% peak-to-valley illumination uniformity across the 3.6 × 3.6 mm^2^ FOV while maintaining compactness (Fig. 1c). Since several neurobiological applications require simultaneous monitoring of neuronal activity for different cell types via dual-color labeling, we designed our miniaturized mesoscope to support this functionality by allowing for the optional integration of two additional LEDs at different wavelengths with an orientation orthogonal to the primary LEDs, together with a dual-passband emission filter (Supplementary Fig. 9a). In both color channels, the dual-color SOMM featured the same 3.6 × 3.6 mm^2^ FOV as the single-color design. Dual-color imaging can be carried out by alternating illumination with either LED color and capturing the respective images in a synchronized manner (Supplementary Fig. 9b).

In miniaturized head-mounted devices that rely on transmission of data and power via cables, the mechanical properties of the cable can strongly affect the animals’ locomotion behavior [20]. To reduce any such detrimental effects, we compared six different multi-core cables with individual core wire diameters ranging from 0.25 to 0.45 mm and found that a minimum core diameter of 0.41 mm was required to transfer the captured images stably (Supplementary Fig. 10a). We then selected the cable with the thinnest jackets available (Supplementary Fig. 10b) and to simplify the assembly, all signal and power cables with the optimized diameter were integrated into a single USB connector (Supplementary Fig. 10c).

In summary, to design SOMM we implemented a principled and comprehensive optimization strategy regarding the above figures of merit, including the form factor, FOV, spatial resolution, excitation efficiency, illumination uniformity, and robustness to axial motion. As a result, SOMM achieved 4 µm lateral resolution across a FOV of 3.6× 3.6 mm^2^ and 300 µm DOF with a device height of only 16.5 mm, enabling cross-cortical imaging of neuronal population activity at cellular resolution at up to 16 Hz. The overall imaging volume (FOV × DOF) accessible by SOMM is 3.9 mm^3^ and thus more than 700 times greater than in the widely used Miniscope v3 [10]. SOMM stands out compared to other miniaturized devices in that it simultaneously meets the requirements of single-cell resolution, mesoscopic FOV, motion robustness, and a low weight compatible with use in freely behaving mice (Supplementary Table 1)

### Performance characteristics of SOMM across a mesoscopic FOV

Guided by the above results and constraints, we designed a mechanical housing for SOMM that maximally reduced weight while maintaining structural stability (Fig. 2a, Data availability). Disk magnets integrated into the SOMM housing as well as the head bars allowed for quick positioning and alignment without the need for anesthesia. The SOMM housing was fabricated using a desktop 3D printer while the DOE was produced using two-photon polymerization (Methods). The assembly of both SOMM and dual-color SOMM is straight-forward and described in detail in open-source documentation (Supplementary Fig. 11, Supplementary Table 2). A custom PCB was inserted into an expansion connector on the back of the sensor PCB to provide power to the LEDs (Supplementary Fig. 10d). To mitigate the extra weight of the additional LEDs in dual-color SOMM, we specially designed a flexible PCB for dual-color SOMM (Supplementary Fig. 9c and 9d) weighing only 0.09 grams, compared to 0.85 grams for a rigid PCB realization. The fully assembled SOMM and dual-color SOMM weighed 2.5 g and 2.9 g, respectively, which represented a weight reduction of >30% compared to other recent mesoscopic head-mounted microscopes [16-18]. The weight difference between SOMM and its dual-color version was negligible and did not affect animal behavior. Specifically, we observed no significant differences in running distance or running speed between mice with SOMM, dual-color SOMM, and control mice inside a 20 cm wide, square, open-field arena, using the optimized cable (Fig. 2b, Supplementary Fig. 12, Supplementary Video 1). Mice carrying SOMM and an un-optimized cable exhibited reduced mobility, highlighting the necessity of a careful optimization of the cable in SOMM (Fig. 2b).

To verify the successful assembly of SOMM, we calibrated the PSF of SOMM with a custom test sample in a parallel manner (Supplementary Fig. 13b and 13c). This photolithographically patterned sample consisted of 1 µm^2^ pinholes separated by 300 µm across a 1 cm^2^ area, facilitating single-shot capture and calibration of PSFs over the entire FOV. As shown in Fig. 2c, the experimentally obtained PSFs at different x, y, and z positions bared resemblance to each other, demonstrating uniform imaging performance across the 3.6 × 3.6 mm^2^ FOV (Supplementary Fig. 13d). Rather than using simulated PSFs, the measured and calibrated PSFs which carry all information regarding fabrication imperfections were used for the deconvolution of our experimental imaging data to ensure optimal reconstruction results. Using the calibrated PSF, we next measured the resolution of SOMM using a customized Siemens Star resolution target across the entire FOV (Fig. 2d). We found that SOMM can faithfully resolve 220 lines/mm after deconvolution across the entire FOV and DOF when using a CMOS sensor with 2.2 µm pixels (Methods). This corresponds to a lateral optical resolution of 4.5 µm and is sufficient to resolve individual neurons.

To confirm that the measured uniformity of resolution was indeed a consequence of our co-optimized DOE design and reconstruction algorithm, we compared SOMM’s performance to that of a miniature imaging device with a conventional spherical lens. To do so, we built a replica of the SOMM housing but replaced our optimized DOE with a plano-convex lens with the same clear aperture as the DOE (Methods). Imaging a single layer of 6-µm fluorescent beads, we found that the images obtained by SOMM exhibited clearly reduced vignetting compared to those captured by the replica system with the plano-convex lens. Quantitatively, SOMM’s illuminated area was ∼3.5 times larger than that of the conventional lens-based replica (evaluated for a relative illumination drop-off threshold of 0.4; Fig. 2e). This allowed SOMM to capture more fluorescent beads at a greater image brightness (Fig. 2f). While the raw SOMM images appeared visually blurrier due to the specific optical encoding performed by the DOE, sharp and crisp images could be restored using our optimized SV-Deconv algorithm (Fig. 2g). At the same time, we validated a key advantage of SOMM when imaging beads close to the edges of the FOV. In these regions, the images captured by the replica with the plano-convex lens were strongly blurred and dim, even after running SV-Deconv. In contrast, the deconvolved images captured by SOMM showed significantly higher intensities and contrast (Fig. 2h). Axially, SOMM captured a larger range of beads than the replica with the plano-convex lens (Fig. 2i, Supplementary Fig. 14a; recorded by imaging a tilted slide carrying fluorescent beads) due to the uniform PSF profiles across the axial dimension (Fig. 2j), resulting in a nearly 6-fold improvement in axial range (Supplementary Fig. 14b). Together with the lateral FOV enhancement, SOMM achieved a 17-fold increase in the accessible and imageable volume.

We next demonstrated that SOMM improves neuron detection sensitivity compared to the replica based on a plano-convex lens. We generated realistically simulated brain volumes with a lateral size of ∼3.6 × 3.6 mm^2^ using NAOMi and simulated the resulting images for both SOMM and the lens-based replica (Supplementary Fig. 15). We found that SOMM achieved remarkably reduced vignetting in capturing brain tissues compared to the lens-based system, because SOMM was thoroughly optimized to eliminate vignetting (Supplementary Fig. 15, Methods). Averaging over n = 5 simulation runs, the neuron density and signal-t0-background ratio (SBR, Methods) delivered by SOMM were both significantly larger than in the lens-based replica (Fig. 2k and 2l).

We further evaluated the robustness of SOMM against axial drifts. We numerically generated fluorescent beads (randomly distributed axially) to approximate neurons displaced by random axial drifting (Supplementary Fig. 7a, b). We note that this is a static approximation of the dynamic axial drifts that can occur during functional recordings. We examine the effect of dynamic drifts directly in the two-photon ground truth experiments described below. In our static simulation studies, we found that SOMM successfully restored compact bead profiles across a 300-µm depth range, with larger values for the SSIM, PSNR, and RMSE metrics compared to the lens-based replica (Supplementary Fig. 7c). To demonstrate the axial robustness in neuronal tissues, we numerically defocused the simulated brain tissues by values ranging from - 150 µm to 150 µm over a FOV area of 3.6 × 3.6 mm^2^, and counted the neurons detected by SOMM and the lens-based replica. We found that compared to the defocus-sensitive lens-based system, SOMM detected many more neurons across all defocusing depths (Fig. 2m). The defocus-induced detection sensitivity loss was also strongly reduced in SOMM compared to the lens-based system, demonstrating SOMM’s superior robustness against axial drifts as they occur during free behavior (Fig. 2m). With the defocus values uniformly distributed in a range of -150 to 150 µm, SOMM detected 1736 ± 123 neurons (mean ± SD, n = 10 simulation runs), a value more than three times larger than the one found for the lens-based system (481 ± 85 neurons; Fig. 2n). SOMM’s significantly greater neuron detection efficiency (p<0.001, two-sided Wilcoxon signed-rank test across, n = 50 random samples and positions) across a large FOV and large range of defocus distances makes it a unique and highly efficient technique for interrogating mesoscopic neuronal population during free behavior.

### Verification of SOMM performance via functional two-photon ground truth imaging

Following the qualitative and quantitative demonstration of SOMM’s imaging performance, next, we showed that SOMM can also accurately capture and demix time-varying neuronal signals *in vivo* as well as perform accurate neuron localization. To achieve this, we designed a computational neuronal processing pipeline for SOMM to effectively eliminate deterioration of temporal and spatial signal quality that arises from crosstalk between neurons, crosstalk between neurons and neuropil, and contamination from fluorescence originating outside of the reconstructed volume. This was done by building a tailored neuronal extraction pipeline that leverages the recently developed DeepWonder technology [39] (Supplementary Fig. 16). DeepWonder is a fast and efficient neuronal extraction technique for widefield fluorescence recordings. Its speed and performance are an order of magnitude improved compared to previous approaches leveraging non-negative matrix factorization techniques [40]. The SOMM data processing pipeline consists of a deconvolution step in which the captured movies are deconvolved using our SV-Deconv algorithm, followed by a segmentation and neuronal footprint extraction step using DeepWonder (Supplementary Fig. 16, Methods). To examine the performance of the proposed processing pipeline, we first tested it on simulated brain tissue data (Supplementary Fig. 17a). We found that the proposed pipeline effectively demixed neurons across a 300 µm axial range (Supplementary Fig. 17b). The mean correlation between ground truth and extracted temporal signals was 0.82 ± 0.18 (mean ± SD, n = 78 neurons), indicating good signal extraction. We further compared the approach to an idealized neuron imaging approach in simulation, in which neurons were sequentially scanned with a diffraction-limited PSF (same NA as SOMM) in 3D. We found no significant differences between these two approaches (Supplementary Fig. 17c, p > 0.1, two-sided Wilcoxon signed-rank test across n = 78 neurons), suggesting that active neurons can be faithfully recorded by SOMM and recovered by our processing pipeline.

To experimentally verify the performance of SOMM in imaging neuron populations, we compared a neuronal time series obtained by SOMM with data obtained using a tabletop one-photon widefield system [41]. In both cases, we imaged the spontaneous activity in head-fixed mice. Although the optical parameters of the tabletop system were partially different from those of SOMM (similar NA of ∼0.1, higher magnification of ∼3x, shallower DOF of ∼40 µm, more sensitive detector response), the statistics of neuronal activities recorded with the two systems can be expected to be similar if SOMM is indeed achieving single-neuron level resolution. We used transgenic animals expressing layer 2/3-specific GCaMP6f calcium indicators for this comparison (Methods). By imaging the same animals alternatingly with either instrument over 3 minutes at 16 Hz, we captured qualitatively comparable images and neuronal calcium activity traces in different locations of the FOV (Fig. 3a). As expected, the histograms of normalized fluorescence changes (ΔF/F) and z-scored extracted neuronal activities acquired across multiple cortical regions showed similar distributions between the two instruments (Fig. 3b and 3c). It is worth noting that in addition to recovering high-quality spatial and temporal neuronal signals like that obtained by the tabletop system, SOMM’s axially uniform PSF results in robustness against axial motion artifacts, which can be significant in freely moving animals.

To perform a direct and quantitative validation of performance and accuracy of SOMM in terms of neuron localization error, fidelity of extracted neuronal signals, and neuron detection performance, we conducted functional ground truth validation with a two-photon (2p) microscope on transgenic animals expressing layer-2/3-specific GCaMP6f calcium indicators (Methods). We coupled our SOMM to a 2p microscope and temporally interleaved SOMM detection under volumetric one-photon illumination with planar 2p imaging, which allowed us to obtain 2p ground truth data at a chosen depth within the SOMM axial range (Supplementary Fig. 18a). Fast alternation between one and two-photon excitation (every 30 ms) allowed us to acquire 2p and SOMM signal nearly simultaneously on the time scale of calcium indicator dynamics. The corresponding ground truth data was obtained by using the well-established CaImAn signal extraction package [30] on the 2p generated data, followed by manual annotation. Using this scheme, due to the use of one-photon volumetric excitation during SOMM acquisitions, the experimental SOMM data directly contained the same out-of-volume fluorescence as is present in standard SOMM acquisitions. Thus, this approach allows for a realistic and quantitative characterization of any detrimental effects due to the out-of-volume fluorescence and crosstalk from axially separated neurons. In addition, we conducted this type of ground truth validation across the entire DOF of SOMM (Supplementary Fig. 18b) and performed *in situ* comparison of SOMM recordings with the temporally interleaved 2p recordings (Fig. 3d).

Comparing this temporally interleaved ground truth dataset to the processed SOMM data, we found that neuron locations and the corresponding time series for the two modalities were highly consistent across conditions where SOMM was focused onto different axial positions (which we refer to as “defocus” for brevity), verifying SOMM’s large DOF (Fig. 3d). A further quantitative examination across the overall 300 µm DOF range corroborated the above results (Supplementary Fig. 19). We found that the mean neuron localization error of SOMM (Fig. 3e) across the defocusing depths of -150 µm to 150 µm DOF was 4.3 ± 2.5 µm laterally (mean ± SD), indicating good neuron localization performance on the scale of the typical neuron size. We compared the extracted temporal neuronal traces with those from the 2p functional ground truth and found a temporal correlation between SOMM and ground truth traces of 0.75 ± 0.19 (median ± SD, n = 439) across all defocus distances (Fig. 3f). To quantify any artifacts introduced by deficient demixing of neuronal signals and suppression of background as a function of spatial separation of neurons, we compared the correlations of neuron pairs found in ground truth as a function of their lateral distances to the correlations of the corresponding pairs found in the SOMM output activity traces (Fig. 3g). We found no significant difference between the correlations in the SOMM-extracted trace pairs and ground truth trace pairs, regardless of the pair distance (Fig. 3g). To investigate the defocus dependence of these pairwise correlations, we computed the excess correlation which was defined as the difference in pairwise correlation between ground truth neuron pairs and the corresponding SOMM neuron pairs (Fig. 3h). We found that at all defocusing depths the modulus of the mean and the standard deviation of the excess correlation values were below 0.07 and 0.25, respectively, indicating robust demixing and discrimination of neuronal signals. To examine the ability of SOMM to find neurons even under defocused conditions, we computed the well-known sensitivity score (true positive rate) for neuron detection between ground truth neurons and SOMM neurons. We found that this score reached values of 0.74 ± 0.15 (mean ± SD) across the examined defocus range (−150 to 150 μm) (Fig. 3i). In summary, the above verifications against 2p functional ground truth demonstrated good neuron extraction performance across the entire DOF range by SOMM.

### SOMM detects thousands of neurons in freely behaving mice at mesoscale

We have shown that SOMM achieves a nearly uniform 4-µm resolution across a 3.6 × 3.6 mm^2^ FOV and across a 300 µm DOF, and corroborated its neuronal detection ability against 2p functional ground truth. We next applied SOMM *in vivo* and demonstrated its capability in recording neuronal population activity across different cortical regions in freely behaving animals (Supplementary Video 2).

We used the capability of our system to record neuronal population activity in freely behaving animals during social interactions with a conspecific (Fig. 4a). Transgenic animals expressing layer-2/3-specific GCaMP6f calcium indicators were used for imaging (Methods). Animals equipped with SOMM were allowed to initially explore an arena before a second mouse of the same sex was introduced, after which recordings were started (Supplementary Video 3). We captured high-contrast activities of 1551 neurons broadly distributed across 11 cortical regions, including the primary somatosensory cortex, primary visual cortex, and retrosplenial cortex during free behavior (Fig. 4b-3d). By tracking the behavior of the animal using DeepLabCut [42], we identified periods of the animals’ interaction with conspecifics and periods without any interactions (Methods). We then investigated the tuning of each neuron to these different states of interaction across tens of cortical regions at high spatial resolution. We found that neurons belonging to the same cortical regions had different tuning properties to social interactions (Fig. 4e), a heterogeneity that can only be captured by studying neural circuits at the single-cell level. Using temporal correlation between neuronal signals as a proxy for functional connectivity, we found that intracortical connectivity within the recorded neuronal population was increased during times when mice were engaged in social behavior: The number of highly correlated neuron pairs (modulus of correlation greater than 0.8) was 263 ± 121 compared to 131 ± 72 (mean ± SD) (Fig. 4f; n = 2 mice), consistent with results from a previous study that used a lower-resolution miniaturized device [17]. These results demonstrated the ability of SOMM to study functional connectivity during free behavior in mice.

Furthermore, we utilized SOMM to study neuronal responses evoked by electrical shocks in transgenic animals expressing layer-2/3-specific GCaMP6f calcium indicators (Methods) in open field arenas; as could also be used in conditioning experiments under naturalistic conditions (Supplementary Video 4). The assay consisted of three stages: during the first stage the animals behaved freely in the open field, in the subsequent stage they received a visual cue followed by electrical shocks from the floor and in the final stage the animals received the same visual cue but no shocks (Fig. 5a). Across an overall 3.8-minute recording, we detected 1314 neurons distributed across 11 cortical regions (Fig. 5b, 5c). We found that the temporal activities of the detected neurons were visibly increased during the second and third stage of the paradigm (Fig. 5d), with a slightly broader distribution of pairwise correlation and lower median correlation values (Fig. 5e). While the animals were relatively stationary before the visual cues and delivery of electrical shocks, locomotor activity was strongly increased and correlated with the evoked neuronal responses demonstrated in the second and third stages of our experimental paradigm (Fig. 5f). The single neuron and high temporal resolution of SOMM allowed us to extract the onset delays of cue-evoked activity peaks in different cortical regions, which were distributed differently across each of the aforementioned stages (Fig. 5g). Spatially, the intracortical connectivity over the same group of neurons was increased in stage two compared to stage one (212 ± 105 and 172 ± 94 highly correlated neuron pairs with modulus of correlation > 0.8, respectively; mean ± SD; n = 3 trials) but reduced in stage three in which animals only received visual cues (130 ± 71 highly correlation neuron pairs; mean ± SD; n =3 trials; Fig. 5h). Jointly analyzing the shock-induced behavior and cortical activities showed that correlations between neuronal activities and motion speeds were significantly increased in stage three compared to stage two (Fig. 5i, ^***^p < 0.001, two-sided Wilcoxon signed-rank test, n = 1314).

Neurovascular coupling plays a critical role in maintaining homeostasis of the cerebral microenvironment [43]. Various microscopy techniques have been developed to study neurovascular coupling [2], but it is challenging to simultaneously observe the activity of neurons and blood vessels in freely behaving mice because this requires the use of dual-color imaging, which in turn adds complexity and weight [44]. Using SOMM, we have overcome this challenge by minimizing device size and weight and have demonstrated dual-color imaging of neurons and blood vasculature without interfering with animals’ behavior (Fig. 2b, Supplementary Fig. 9, 12). We thus conducted dual-color SOMM recording using tetramethylrhodamine dextran (TRITC-dextran)-labeled blood vessels and GCaMP-labelled neurons while recording the fluorescence of the two different indicators by temporally interleaving the two channels (Fig. 6a, Methods). During a 4-minute recording session, individual neurons across the motor, visual, and retrosplenial cortices were faithfully identified and tracked (Fig. 6b) along with rich vascular networks (Fig. 6c), both imaged at 8 Hz (Methods). Dual-color SOMM thereby opens the possibility to study neurovascular regulation across cortical areas in walking and running animals. To illustrate this, we summarized the activity of neuronal populations by principal component analysis (PCA) and studied the correlation between the principal components (PCs) and vascular dynamics as represented by changes in fluorescence due to modulation of blood flow. We found that the first neuronal PC exhibited low and along the posterior–anterior axis a spatially homogeneous correlation with the vascular dynamics, while the second and third neuronal PCs showed a positive correlation on the anterior side and a negative correlation on the posterior side with the vascular dynamics (Fig. 6d). Such correlational observations between large-scale neuro and vascular dynamics during free behavior are only possible thanks to SOMM’s mesoscale FOV and simultaneous dual-color imaging capability and to our knowledge is not possible with any system to date.

## Discussion

In summary, we present a new principled design approach and implementation of an optimized head-mounted miniaturized mesoscope. Our SOMM system enables large-scale cellular resolution neuronal population imaging, offers dual-color imaging capability, and robustness against axial motion-induced drifts. By systematically optimizing the imaging path, illumination path and reconstruction algorithm jointly and iteratively (while minimizing weight and optimizing the device form factor), SOMM achieves 4-µm optical resolution across a 3.6 × 3.6 mm^2^ FOV and 300 µm DOF at a weight of less than 2.5 grams. Equipped with LEDs containing different colors and flexible PCBs, the dual-color SOMM can image two different fluorescent labels without crosstalk.

While there are well-established procedures and realizations for optimizing specific microscope parameters such as its resolution [10] or FOV [17], our demonstrated design approach for SOMM is based on a *principled* and comprehensive optimization of multiple parameters simultaneously with the aim of allowing optimal population-level *in-vivo* neuronal recordings. SOMM establishes a simple and extensible platform that can be customized and adapted to other mammalian model systems and opens up the door to a better understanding of the neuronal basis of social interactions. In particular, future neuronal dynamics underlaying interactions of multiple animals can be studied by integration of available wireless data transmission technology into our current system [16] allowing for fully unconstrained animal behavior.

Given the compact footprint of SOMM afforded by the use of diffractive optics, it is also possible to position multiple SOMM devices on different cortical areas for assessing different brain structures simultaneously [19] and deeper cortical regions can be accessed by combining microprisms that transform axial side views into laterally oriented planes [45], GRIN lenses, or glass pillar periscopes with SOMM [41, 46]. Considering that the size of the FOV offered by implantable optics is usually smaller than that of SOMM, these arrangements will also allow for simultaneous recording of neuroactivity of select regions at depth together with the activity of larger more superficial brain areas.

By combining the flexibility of diffractive optics with a powerful optimization strategy, our approach offers a platform suitable for a broader range of other imaging applications such as air quality monitoring [47] and parasite screening [48], where a combination of large FOV, reduced aberrations under mechanical and form constraints are required. As such we anticipate that the proposed method can serve as a guideline for optimizing integrated optical observation devices.

## METHODS

### Systematic optimization scheme

*Form factor optimization*. In the first round of optimization, we used first-order optics to characterize the sample distance *d*_1_, sensor distance *d*_*2*_, diffractive optical element (DOE) size *L*, sampling ratio, field-of-view (FOV), and system thickness. We aimed for single-neuron resolution, which means that the diffraction-limited spot size should be at least 2 times smaller than the neuron size under Nyquist sampling criteria, and the diffraction-limited spot must be sampled by at least 2 pixels. These requirements can be written as

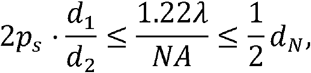

with the numerical aperture *NA =* sin(atan(*L*/2*d*_*1*_)) ≈ *L*/2*d*_*1*_ and the neuron size *d*_*N*_, which is roughly 10∼12 µm in the mouse brain[37]. Note that although the DOE’s phase *ϕ* will alter the spots shape, the minimum feature of the PSF will still be determined by the Rayleigh criterion, 1.22 *λ* / *NA*, and needs to be sampled by at least 2 pixels to be resolved by deconvolution. For the off-the-shelf sensors used in common miniaturized microscopes, usually pixel size *p*_*s*_ ≥ 2.2 µm and active area *D* ≤ 5 mm. The maximum achievable FOV in the single-lens imaging system thereby can be determined by FOV *= D · d*_1_ / *d*_2_ On the other hand, the angular response curves of the sensor pixels will clip rays incident at an angle greater than a threshold *α* [34], which means the FOV is determined by

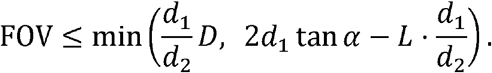

Usually, *α* will be 20°∼30° for a standard array sensor [34]. Further considering that the DOE is relatively flat [50], the thickness of the device can be approximated as *d*_*1*_ *+ d*_*2*_

To optimize the optical system for mesoscale imaging in a lightweight and small volume, we ran a brute-force search for maximizing FOV under the above constraints. The optimization proceeded with three summarized variables *β *=* d*_1_ / *d*_*2*_, *L*, and T *= d*_1_ + *d*_*2*_ These constraints shaped the potential solution into a manifold (Fig. 1a), and we labeled the FOV values with different colors. We found the largest FOV under these conditions was ∼4.9 mm laterally with a total thickness ∼23 mm. Such an optimal condition (marked by a hollow star in Fig. 1a) was achieved by *d*_1_ ≈ *d*_*2*_ and *L* ≈ 2.9mm. Such a thickness, however, would be too large because the corresponding device weight would prevent unrestrained movement. On the other hand, we observed that along the black line in Fig. 1a the FOV did not change much. With the thickness in mind, we thus choose *L* ≈ 1.5 mm along that line (marked by a solid star), which gives a maximum FOV of 4.5 mm with thickness T ≈ 14 mm and β ≈ 1. The thickness of this design is 40% less than the popular Miniscope v3 system without the sensor[10], with a FOV increased by 50 times.

#### Diffractive optical element (DOE) optimization

The imaging model in the above form factor optimization was ideal, i.e., aberrations, noise, and limited contrast were not considered. To translate the optimized framework into a practical system, we used prior information on neuron sizes to optimize a versatile phase element such that it provided uniform imaging resolution across a large volume. Let us assume a neuron under the dura is located at a distance *d*_*1*_ *+* Δ*z* from the DOE, with the DOE a distance *d*_*2*_ away from the active area of the sensor (Fig. 1b). If we take a neuron as a point source at (*x*_0_,*y*_0_) the light field that reaches the DOE is

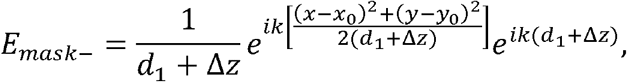

If we assume that the DOE has a square aperture with a phase shift *ϕ* (*x*_1_,*y*_1_), the DOE-encoded light field is then

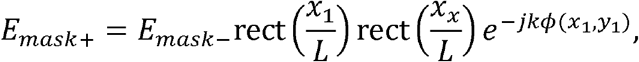

where *L* is the DOE size. The height map Δ of the DOE is related to the phase shift *ϕ* via Δ *= ϕ /(n − 1*), and *n* is the refractive index of the DOE material. The DOE-encoded light field further propagates to the sensor as

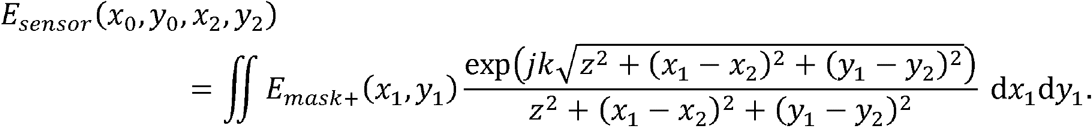

The final PSF from the fluorescent neuron is then *I*_*PSF*_. *=* | *E*_*sensor*_ |^2^ The reason that we used a non-approximated angular spectrum method instead of Fresnel propagation as in previous work [35] was that the desired FOV in our system was beyond the paraxial approximation. The final captured sensor image is

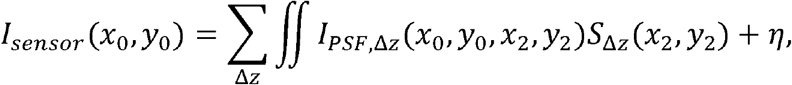

where *I*_*PSF,Δz*_ and *S* _Δz_represented PSFs and objects from different depths Δ z *η* ∼ *N* (0,*σ*^*2*^) is additive noise in the sensor. For a standard imaging system (i.e.,*ϕ* implements a spherical lens), *I*_*PSF*_ becomes blurred as the object moves away from the focal plane due to motion or as it moves toward the FOV boundaries due to coma; causing non-uniform imaging quality across the target volume. To overcome these degradations, we used a loss function to measure the uniformity of the PSF by evaluating the similarity of reconstruction across the whole volume (Supplementary Fig. 1). The advantage of this approach compared to directly measuring the similarity of PSFs was that it focused on the quality of the final output of the system, which meant the optimization would directly improve the final output quality. Further, we added energy constraint loss around the centroid of the PSF to make sure that it had high contrast for high SNR. The loss function that was used to optimize *ϕ* could be written as

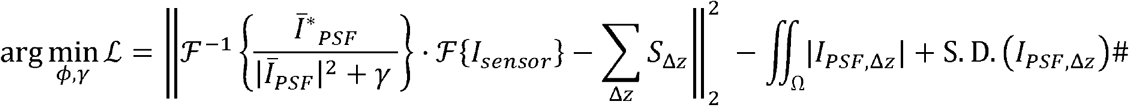

where *γ* is a regularizer, *Ī*_*PSF*_ is the Fourier transform of the PSF *I*_*PSF*_ at z *=* 0, * indicates complex conjugation, Ω is a small window around the centroid of the PSF and S.D is the standard deviation operator. The first part of the loss function compares information recovered via the well-known Tikhonov deconvolution from the 2D capture *I*_*sensor*_ with the ground truth Σ_Δz_ *S*_Δz_The second term encourages high PSF contrast by ensuring that photons from a single neuron do not spread out too much; introducing this term lowers the signal-to-noise ratio of the final capture. The third part forces the PSF at different positions to have the same peak intensity such that the imaging system exhibits minimum vignetting and large DOF.

We expand the optimization variable *ϕ* into a sum of a basic lens and a set of Zernike basis functions,

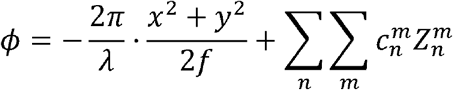

where

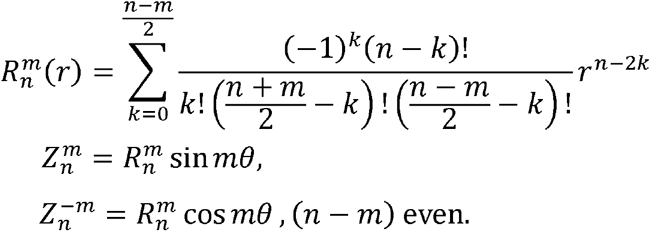

We only optimized the corresponding Zernike coefficients while ignoring the constant, tip/tilt, and defocus terms in the Zernike bases. The choice of optimizing a Zernike polynomial-based model instead of a pixel-wise model had two advantages. First, the parameter number that needed to be optimized was reduced by a factor of 2,500 in the Zernike method compared to the pixel-wise optimization. Second, the Zernike method satisfied the bandwidth limitation of the angular spectrum method in numerical simulation, which produced more reliable results (Supplementary Note 1).

#### Illumination optimization

For this approach, we conducted illumination optimization with LightTools (Synopsys), exported a 3D model of SOMM from SolidWorks (SolidWorks 2019; Dassault Systèmes) and imported the model into LightTools. We also created a virtual LED source and simulated illumination rays along the illumination tubes inside the SOMM body using a Monte-Carlo approach and placed virtual detectors in the focal plane. In turn, the detected ray density at different field positions was set as a merit function. Using greedy search, we tested different LED distances and illumination angles and chose the most uniform configuration as the final setup for SOMM. The details of the illumination paths can be found in the open-sourced SOMM mechanical design files (Data availability).

#### Realistic brain tissue simulation

To synthesize realistic cortical tissue and generate the corresponding widefield capture, we used the Neural Anatomy and Optical Microscopy (NAOMi)[37] package. By implementing this package, a brain tissue volume was populated with multiple blood vessels, as well as neuron somata, axons, and dendrites. Neurons and dendrites were assigned synthesized fluorescence activity that reflected their calcium dynamics. While NAOMi was originally designed to simulate two-photon excitation, we modified the original NAOMi pipeline such that it could faithfully simulate data acquisition in one-photon imaging. We changed the excitation wavelength from the near-infrared range into the visible range and added tissue-induced aberrations into DOE-generated PSFs to resemble the potential wavefront distortions in in vivo imaging.

To ensure that the DOE in SOMM achieves robust performance across a large FOV (3.6 × 3.6 mm^2^) and a large DOF (300 µm), the NAOMi-generated tissue volumes (200 × 200 × 60 µm^3^) were randomly translated laterally and axially to mimic the effects of axial drifts during the DOE optimization (Supplementary Fig. 1b).

#### Shift-variant deconvolution algorithm

For the output of the DOE optimization step, we achieved both an optimized PSF *I*_*PSF*_ and an optimized deconvolution *λ* parameter ; both of which are sensitive to the position in the FOV. As a result, we tiled the captured raw image into patches to conduct patch-wise deconvolution (i.e., shift-variant deconvolution). For a raw patch *I*_*raw*_,(*x*_0_,*y*_0_)centered at (*x*_0_,*y*_0_), the deconvolved patch is

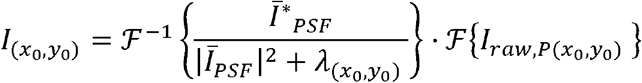

where *λ*(*x*_0_,*y*_0_) is the optimized deconvolution parameter at (*x*_0_,*y*_0_), *Ī*_*PSF*_ is the Fourier transform of the optimized PSF *I*_*PSF*_ at z *=* 0 and (*x*_0_,*y*_0_), and * indicates complex conjugation. We conducted the above deconvolution for each patch, stitched those patches into a final output image and cropped out overlapping areas between patches. We termed the above procedure as the SV-Deconv algorithm. An additional realization was that the SV-Deconv algorithm could be accelerated by being implemented for GPUs. Since there are inevitable differences between designed DOEs and fabricated DOEs, we conducted SV-Deconv on the experimental data based on experimentally calibrated PSFs (Supplementary 13) but by utilizing the deconvolution parameter *λ* from the optimization stage.

#### Fabrication and assembly details

*DOE* fabrication. The DOE was fabricated using two-photon photolithography (service provided by Moveon Technologies, using a Nanoscribe Professional GT system) in the high-resolution dip-in liquid lithography (DiLL) configuration on a 170-μm-thick fused silica substrate. The fabricated DOE had an overall height of 1 μm with a 150 nm height step size, and a lateral pixel size of 2 μm. After photolithography, the substrate was cut into 1.5 × 1.5 mm for assembly.

#### Housing fabrication

SOMM was designed using a CAD program (SolidWorks 2019; Dassault Systèmes) and the full design files are available in the open-sourced SOMM mechanical design files (Data availability). The top and bottom housing were 3D-printed using a desktop stereolithography printer (microarch P140, BMF Precision Tech Inc.) with black poly (methyl methacrylate) (PMMA) resin. Two blue LEDs wired in series using 26-gauge wires (LUXEON Rebel Color Blue, 470 nm; Digi-Key) and a custom-diced band-pass excitation filter (450–490 nm, 4 × 4 × 1 mm^3^, ET470/40x; Chroma) were installed into the illumination slot and fixed by ultraviolet (UV)-curable optical glue (AA352 Light Cure Adhesive; LOCTITE) (Supplementary Fig. 11). The fabricated DOE was gently inserted, press-fitted into a 3D-printed carrier and glued in place. The carrier was then inserted into a rectangular slot at the top side of the bottom housing, followed by another custom-diced band-pass emission filter (500–550 nm, 4 × 4 × 1 mm^3^, ET525/50m; Chroma) (Supplementary Fig. 11). A CMOS sensor (MT9P031, Aptina) was glued to the top housing. This sensor was used with a readout ROI matched to the FOV boundary to increase the frame rate. The top housing was slid onto the rectangular shaft of the bottom housing; the focus was adjusted and then fixed using stainless steel 0–80 screws. Two circular neodymium magnets (diameter 3 mm) were bonded to the bottom surface of the bottom housing with cyanoacrylate glue (Supplementary Fig. 11). For dual-color SOMM, we used a bandpass filter (ET470/40x, Chroma) for blue channel excitation, and another bandpass filter (ET570/20x, Chroma) for lime channel excitation. We used a dual-pass filter (59022, Chroma) as an emission filter to select both green and red fluorescent photons. The excitation and emission filters were selected to avoid leakage of excitation light to the sensor (Supplementary Fig. 9).

#### SOMM replica with a plano-convex lens

To compare the proposed DOE in SOMM with a plano-convex lens head-to-head, we swapped the DOE in SOMM for a plano-convex lens (Edmund 65-287). The selected lens had a similar effective focal length (4 mm) as the fabricated DOE (3.5 mm) and shared the same aperture window (1.5 mm in diameter), detection filters, and sensors as the DOE. The internal illumination and detection tubes of the replica were similar to those found in SOMM as well.

#### Wiring

A customized cable was used to connect the CMOS sensor to the PC (Supplementary Fig. 9, 10). For dual-color imaging, LED control signals were sent from a microcontroller (Arduino Uno; Digi-Key) to drive alternating illumination with blue and lime light. Two separate power sources (3.3V tunable sources) were used to modulate the current delivered to the blue and lime LEDs. The microcontroller also sent an external trigger input to the CMOS sensor for synchronizing illumination and frame capture. All captured frames were transmitted to the PC over a USB 2.0 connection.

#### Hybrid “temporally interleaved” 2p–SOMM functional ground truth recordings

To validate SOMM, we modified both the illumination and detection arms of a tabletop two-photon microscope (Supplementary Fig. 18a). A titanium-sapphire laser system (MaiTai HP, Spectra-Physics) served as the two-photon excitation source (920 nm central wavelength, pulse width <100 fs, 80 MHz repetition rate). A half-wave plate (AQWP10M-980, Thorlabs) and an electro-optic modulator (350-80LA-02, Conoptics) were used to modulate the excitation power. A 4f relay system (AC508-200-B and AC508-400-B, Thorlabs) with a 2× magnification was used to expand the laser beam and relay it onto a resonant scanner (8315K/CRS8K, Cambridge Technology). The scanned beam went through a scan lens (SL50-2P2, Thorlabs) and a tube lens (TTL200MP, Thorlabs) and was focused by a high numerical aperture (NA) water immersion objective (20x/0.4 NA, MY20X-824, Mitutoyo). A high-precision piezo actuator (P-725, Physik Instrumente) moved the objective for fast axial scanning. A long-pass dichroic mirror (DMLP650L, Thorlabs) was used to separate fluorescence signals from the femtosecond laser beam by reflecting the fluorescence signals and transmitting the infrared laser light. The back aperture of the objective was optically conjugated to the detection surface of the PMT with a 4f system (TTL200-A and AC254-050-A, Thorlabs).

For the SOMM excitation path, a long-pass dichroic (DMLP505L, Thorlabs) in the original detection path of 2p microscope was used to send blue light from an LED (M470L4-C1 and MF475-35, Thorlabs) to the objective. For the SOMM detection path, we used a second identical pair of objectives (20x/0.4 NA, MY20X-824, Mitutoyo) and tube lens (TTL200MP, Thorlabs) to relay the widefield-excited fluorescent light field from the first (main) objective and tube lens to the detection plane of SOMM. The entirely identical magnification module (the first group of the objective and the tube lens) and demagnification module (the second group of the objective and the tube lens) delivered the fluorescent light field with minimized optical aberrations and uniform magnification. A 50:50 (reflectance : transmission) non-polarizing plate beam splitter (BSW27, Thorlabs) was placed after the widefield dichroic (DMLP505L, Thorlabs) to split fluorescent signals for PMT-(PMT1001, Thorlabs) and SOMM-detection, respectively. Fluorescence filters (MF525-39, Thorlabs; ET510/80M, Chroma) were placed in front of both the PMT and SOMM to fully block both the femtosecond laser and widefield excitation beam.

To avoid excitation crosstalk and protect the PMT from high-flux widefield emission photons, we added a linear galvo that served as an optical shutter for the PMT detection path by deflecting widefield fluorescent photons when the LED was on [39]. We further configured the EOM to block laser light during widefield imaging. The LED (M470L4-C1) was used in external trigger mode with a typical rise and fall time of less than 1 ms, with reduced on-time intervals relative to the frame rate to avoid PMT overexposure.

To introduce an offset between the 2p focal plane and the SOMM focal plane for investigating SOMM performance during defocusing, the entire SOMM body was translated with a linear translation stage (XR25P/M, Thorlabs). Both modalities used frame rates of 15 Hz, and defocus configurations ranging from -150 to 150 μm at ∼100 μm depth in the mouse cortex.

#### Experimental model and subject details

Adult mice (male or female without randomization or blinding) at 8–16 postnatal weeks were housed in an animal facility (24 °C, 50% humidity) under a reversed light cycle in groups of 1–5.

For functional imaging of neural activity in comparison with tabletop one-photon widefield system, two-photon microscope imaging, as well as studying social interactions and fear conditioning, we used transgenic mice hybridized between Rasgrf2-2A-dCre mice and Ai148 (TIT2L-GC6f-ICL-tTA2)-D mice expressing Cre-dependent GCaMP6f genetically encoded calcium indicator (GECI). Briefly, mice were first anesthetized with 1.5% (by volume in O2) isoflurane and a 4.0-mm diameter craniotomy was made with a skull drill. After removing the skull piece, a coverslip was implanted on the craniotomy region, and a titanium headpost (see Data Availability) was then cemented to the skull for the SOMM mounting. After the surgery, 0.25 mg/g (body weight) trimethoprim (TMP) was injected intraperitoneally to induce the expression of GCaMP6f in layer 2/3 cortical neurons across the whole brain. Mice were subsequently allowed to recover for 14 days before the imaging experiments.

For neurovascular imaging with dual-color labeling, 100 μl solutions containing 50 mg/ml of 2000 kDa TRITC-Dextran in PBS (Sigma or Thermo-Fisher Scientific) were injected into the circulation of the transgenic animals via the tail vein cannula.

#### Freely moving experiments

##### Arena

We placed mice into a homemade arena consisting of a plastic box (∼200 × 200 × 200 mm^3^ size), an IR-sensitive camera (BFS-U3-51S5M-C, Point Grey), and a 940-nm IR lamp.

##### Focus adjustment of SOMM

Before every experiment, mice were lightly anesthetized (0.5–1% isoflurane in pure oxygen) and head-fixed in a stereotax to clean the cranial window of any debris. The SOMM device was then securely mounted on the implant. The blue LED was switched on and the position of the top housing relative to the bottom housing was manually adjusted until the neurons were in focus and the locking screw was tightened to secure the position. Once focused, the mice were released into the center of the open field arena and were ready for experiments.

##### Open field experiments

Mice underwent acclimatization for 3–5 days during which an experimenter handled each mouse for 5–15 minutes. Mice were fitted with a SOMM replica weighing the same as a fully assembled SOMM during the handling period. For experimentation, the SOMM was fitted onto the mouse and the mouse was quickly transferred to an open field arena. Experimental trials lasted 6 minutes, including a 3-minute warm-up phase.

##### Social behavior experiments

During the social behavior experiments, a C57BL/6 mouse of the same sex as the imaged mouse was gently introduced into the arena by an experimenter 5 minutes after the initiation of the trial.

##### Conditioning experiments

During the conditioning experiments, the SOMM was fitted onto the mouse and the mouse was quickly transferred to an open field arena. The animal first freely explored the arena for ∼4 minutes (stage 1). Then, the animal received periodic visual cues (lasting 2 seconds, from a 650 nm light bulb) and followed by electrical foot shocks (duration 0.5 seconds, 0.75 mA) for 1 minute (stage 2). Finally (stage 3), the animal received only visual cues for 1.5 minutes (duration 2 seconds, from a 650 nm light bulb).

### Data analysis

#### Behavior video analysis

Videos of mouse behavior were captured in AVI format using an overhead camera (BFS-U3-51S5M-C, Point Grey). The mice were tracked using DeepLabCut [42]. Tracking output from each trial was manually verified to ensure tracking accuracy. A 20-cm square area was defined at the center of the arena as the ‘open field’. For social interaction experiments, we manually scored behavior with 1-second precision based on whether the mouse with the SOMM was interacting with the other mice or not.

#### Time-lapse neuronal data processing

Captured raw data from SOMM was first frame-by-frame deconvolved with the proposed algorithm (Code Availability), and then motion-corrected using the open-source NormCorre algorithm in non-rigid mode [51]. The motion-corrected movies were processed by the recently developed DeepWonder [39] package to extract reliable neuronal segments and signals. DeepWonder was fine-tuned to the detailed characteristics of the SOMM modality following the guidelines in the DeepWonder manuscript [39]. The Allen Brain Atlas was used as an anatomical reference for data analysis in this study (http://www.brain-map.org) [49].

### Performance metrics

#### Cross-coherence

The cross-coherence between any two PSFs is defined as ‖*PSF*_1_ * *PSF*_*2*_ ‖_∞_ *=* max [*PSF*_1_ * *PSF*_*2*_ ]where * represents the 2D correlation and max[·] is the element-wise maximum. Higher values of cross-coherence represent the greater similarity between *PSF*_1_ and *PSF*_*2*_

#### Optical transfer modulation

A customized resolution chart was used to model the expected optical transfer through the system at the center of the imaging plane (the optical axis) and 1.5 millimeters lateral to the center (Fig. 2d). The transfer function is expressed as the normalized amplitude modulation of gratings with varying spatial frequencies (line pairs/mm) averaged over all directions as *M =* (*I*_*max*_ − *I*_*min*_)/ (*I*_*max*_ + *I*_*min*_)where *I*_*max*_ and *I*_*min*_ are the pixel values of the light and dark stripes.

#### Signal-to-background ratio (SBR) calculation

We calculated SBR through the following steps: First, we segmented the neurons by template matching, then we averaged the signal from the neuron area and averaged over a ring surrounding the neuron to estimate the background signal. SBR was then taken to be the ratio of the signal and background values.

#### Structure similarity (SSIM)

Structural similarity index (SSIM) is a widely used metric for the assessment of visual quality of images and remote sensing data. We used the *ssim()* function in MATLAB to calculate the similarity between PSFs and the similarity between reconstructed images and ground-truth images.

#### Peak-signal-to-noise-ratio (PSNR)

Peak signal-to-noise ratio (PSNR) is an engineering term for the ratio between the maximum possible power of a signal and the power of corrupting noise that affects the fidelity of its representation. We used the *psnr()* function in MATLAB to calculate the similarity between reconstructed images and ground-truth images.

## DATA AVAILABILITY

Example data as well as all CAD files for manufacturing the SOMM are available at https://github.com/yuanlong-o/SOMM. The raw and analyzed datasets generated during the study are too large to be publicly shared, but they are available for research purposes from the corresponding authors upon reasonable request.

## CODE AVAILABILITY

The custom code that comprises the SOMM optimization and calcium analysis pipeline is available in Supplementary Software. Future updates to the code will be published at https://github.com/yuanlong-o/SOMM, together with optical and optomechanical designs, optimization code used for designing SOMM, and SOMM video processing code.

## ACKNOWLEDGMENTS

We thank Qiyu Zhu for providing dual-color labeled GCaMP mice and Peer Strogies and James Petrillo at The Rockefeller University’s Precision Instrumentation Technology (PIT) for fabrication of mechanical components. This work was supported by the National Natural Science Foundation of China (No. 62088102), the Ministry of Science and Technology of the People’s Republic of China (No. 2020AA0105500), the National Institute of Neurological Disorders and Stroke of the National Institutes of Health under award numbers 1RF1NS110501 (A.V.) and 1RF1NS113251 (A.V.), the Kavli Foundation through the Kavli Neural System Institute (A.V.) and through a Kavli Neural Systems Institute postdoctoral fellowship (T.N.)

## AUTHOR CONTRIBUTIONS

Y.Z. designed and implemented the SOMM optimization pipeline, performed simulations, designed the optics and mechanics in SOMM, contributed to SOMM assembly, and wrote the manuscript. L.Y. contributed to SOMM assembly and calibration experiments, performed freely behaving experiments, analyzed data, and wrote the manuscript. J.W. provided critical support on system setup and signal processing and wrote the manuscript. T.N. provided important suggestions, critical support on system setup and contributed to manuscript editing. R.Z. and G.X. performed cranial window surgeries, viral injections and contributed to freely behaving experiments. M.W. and H.X. provided critical support on system setup. A.V. conceived and led project, conceptualized and guided Y.Z. on the implementation of the principled optimization approach, designed experiments, guided data collection and analysis and wrote the manuscript. Q.D. led and co-funded the project, co-supervised Y.Z., supervised L.Y., J.W., R.Z., G.X., M.W. and H.X. and wrote the manuscript.

