## Supplemental Information for "A Systematically Optimized Miniaturized Mesoscope (SOMM) for large-scale calcium imaging in freely moving mice"

**Supplementary Figure and Table Titles**

| Supplementary Figure 1 | Realistic training data for SOMM generated by NAOMi. |
| --- | --- |
| Supplementary Figure 2 | Cross-coherence metric increasing during the DOE optimization stage of SOMM, across various lateral and axial positions. |
| Supplementary Figure 3 | Characteristics of the optimized DOE in SOMM compared to a plano-convex lens. |
| Supplementary Figure 4 | Comparison of proposed SV-Deconv algorithm with standard, shift-invariant deconvolution applied to a large FOV. |
| Supplementary Figure 5 | Resolution of SOMM evaluated by numerical simulation. |
| Supplementary Figure 6 | Specifications of a system of spherical lenses with the same optical resolution and FOV as SOMM. |
| Supplementary Figure 7 | Large depth of focus due to elongated PSF in SOMM. |
| Supplementary Figure 8 | Effects of imaging through a dichroic in a single-lens imaging system akin to SOMM. |
| Supplementary Figure 9 | Dual-color SOMM enables dual-color imaging in a compact form. |
| Supplementary Figure 10 | Optimization of cables in SOMM. |
| Supplementary Figure 11 | Step-by-step assembly illustration for SOMM and dual-color SOMM. |
| Supplementary Figure 12 | Negligible effects of carrying SOMM and dual-color SOMM on mouse motion statistics. |
| Supplementary Figure 13 | PSF calibration for SOMM using a customized pinhole array. |
| Supplementary Figure 14 | Characteristics of SOMM compared to a SOMM replica with a plano-convex lens when imaging axially tilted samples. |
| Supplementary Figure 15 | Characteristics of SOMM in simulated neuronal recordings compared to a lens-based SOMM replica. |
| Supplementary Figure 16 | Calcium imaging processing pipeline of SOMM. |
| Supplementary Figure 17 | Validation of calcium signal extractions in SOMM. |
| Supplementary Figure 18 | System setup for temporally interleaved 2p–SOMM functional ground truth recordings. |
| Supplementary Figure 19 | Temporally interleaved 2p–SOMM functional ground truth validation. |
| Supplementary Table 1 | Comparison of SOMM and other miniaturized microscope. |
| Supplementary Table 2 | Component parts list of SOMM. |
| Supplementary Note 1 | Details of end-to-end DOE optimization. |

**Supplementary Video Captions**

| Supplementary Video 1 | Evaluation of the animal movements when wearing SOMM and dual-color SOMM in an open field arena (size 200 × 200 × 200  mm^3^). First column: animal with a headbar only. Second column: Same animal wearing SOMM, connected with the optimized cable. Third column: Same animal wearing dual-color SOMM, connected with the optimized cable. Fourth column: Same animal wearing SOMM connected with unoptimized wires. Top row: raw videos. Bottom row: Extracted animal motion tracks. |
| --- | --- |
| Supplementary Video 2 | SOMM enables mesoscopic observation of cortical activities during free exploration of an open field arena by solitary animals. Left column: raw behavior video recordings (top) and extracted motion traces (bottom). Middle column: Heatmap of extracted calcium activities of 877 neurons (rows), shown for a sliding window centered on the current timepoint. Neurons sorted by onset time of largest activity peak. Right column: raw (top) and background-removed SOMM neuronal recordings. Scale bar: 500 μm. |
| Supplementary Video 3 | SOMM enables mesoscopic observation of cortical activities during animal interactions. Left column: raw behavior video recordings (top) and extracted motion traces (bottom). The two animals are represented as red and blue lines, respectively. Increased line thickness indicates contact between animals. Middle column: calcium activities of 1072 neurons (rows), shown for a sliding window centered on the current timepoint. Neurons sorted by onset time of largest activity peak. Right column: raw (top) and background removed (bottom) SOMM neuronal recordings. Scale bar: 500 μm. |
| Supplementary Video 4 | SOMM enables mesoscopic observation of cortical activities during free behavior in an open field arena under electric foot shocks. Left column: raw behavior video recordings (top) and synchronized visualization of current experimental condition (bottom): cue (blue), shock (yellow), and trial intervals (gray). Middle column: calcium activities of 1003 neurons (rows), shown for a sliding window centered on the current timepoint. Neurons sorted by onset time of largest activity peak. Right column: raw (top) and background removed (bottom) SOMM neuronal recordings. Scale bar: 500 μm. |

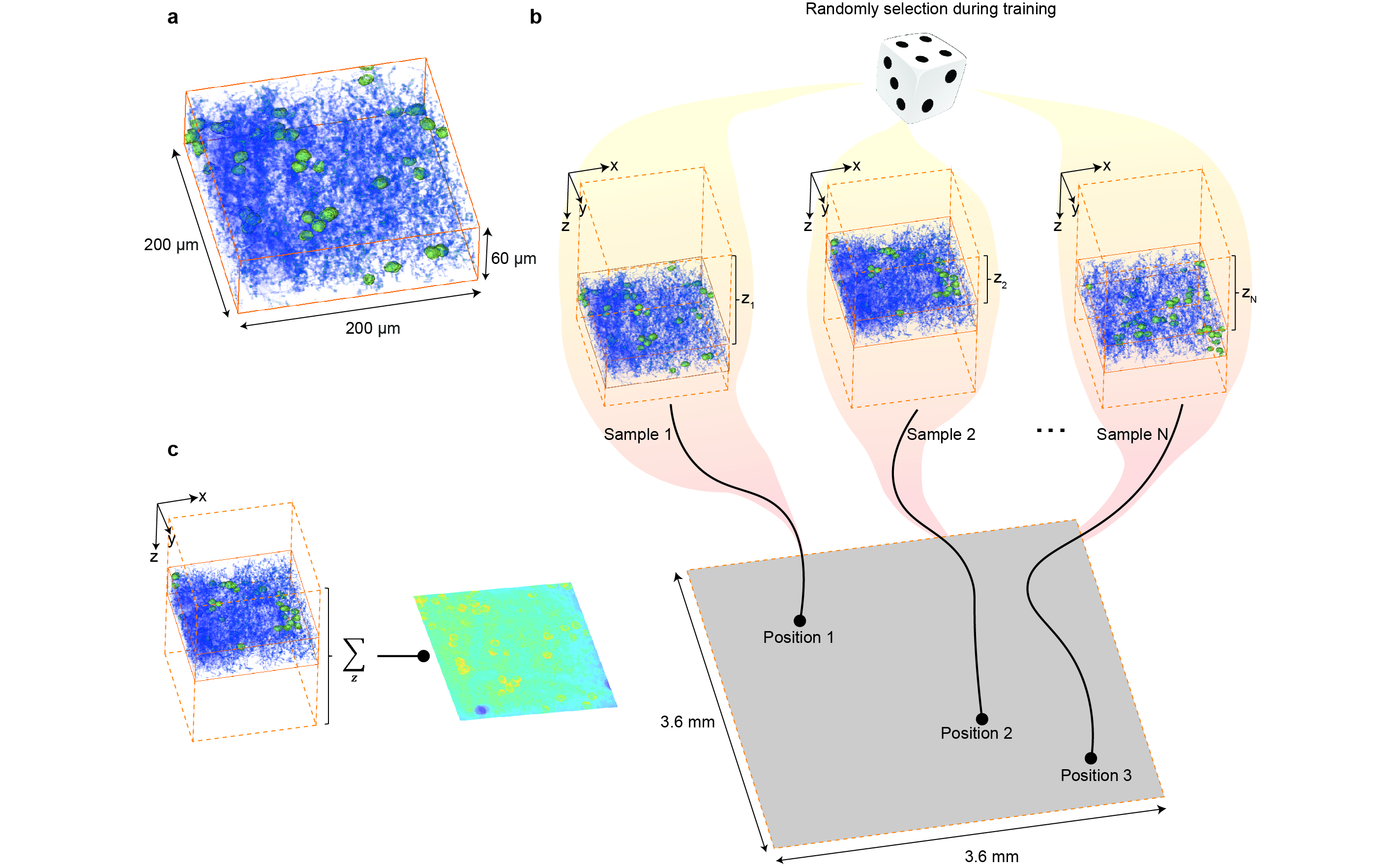

Supplementary Figure 1

Realistic training data for SOMM generated by NAOMi.

**a**. Simulated volume of brain tissue generated using the NAOMi package [1]. The simulated cortical tissue had a thickness of 60 µm, which corresponded to the 1/e^2^ width of an ideal PSF with the same NA as SOMM (NA 0.1).

**b**. Illustration of training randomization strategy. During the optimization of the DOE in SOMM, random training data samples were selected, and shifted into different axial positions (z_1_, z_2­_, …, z_N_, varying across 300 µm range) and lateral positions (Position 1, Position 2, …, Position 3, varying across 3.6 × 3.6 mm range) before being convolved with the optical PSFs. By virtue of these randomizations, the optimized DOE in SOMM delivered consistent performance across a 3.6 × 3.6 mm field of view (FOV) and 300 µm depth of field (DOF).

**c**. Illustration of ground truth generation for training. The NAOMi-generated volume was shifted randomly along the axial direction, and then summed along z axis to ensure that the DOE optimization results in a DOE phase pattern that is insensitive to axial drift.

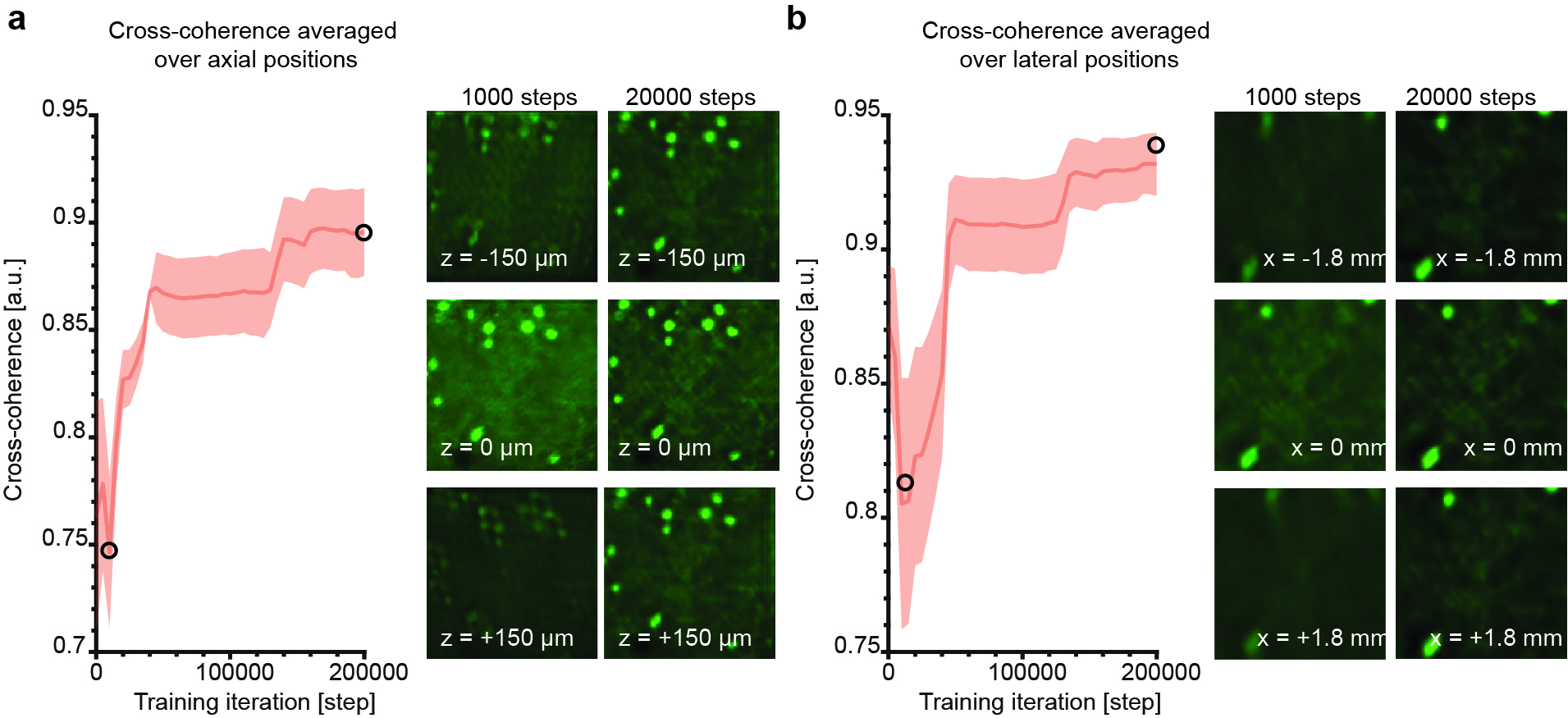

Supplementary Figure 2

Cross-coherence metric increasing during the DOE optimization stage of SOMM, across various lateral and axial positions.

**a**. Left: Cross-coherence of reconstructed training data with ground truth, averaged across a range of depths (solid line: mean, shaded area: SD), showing an increase with training iterations. Right: comparison of reconstructed test data (i.e., held out from training) for the DOE generated after 1,000 training iterations (left column), and after 20,000 training iterations (right column), for different axial depths. After 1,000 training iterations, the reconstructed test data frame in the focal plane was clear and exhibited high contrast, while at large distance (defocus) from the focal plane (z = -150 µm and z = 150 µm), the reconstruction quality was degraded, with comparably low cross-coherence with ground truth (not shown). After 20,000 training steps, the cross-coherence with ground truth in both the focal plane and at defocused depths was greatly improved.

**b**. Left: Cross-coherence of reconstructed training data with ground truth averaged over different lateral positions (solid line: mean, shaded area: SD). Right: comparison of reconstructed test data (i.e., held out from training) for the DOE generated after 1,000 training iterations (left column), and after 20,000 training iterations (right column), for different lateral positions. After 1,000 training iterations, the reconstructed test frame in the center of FOV was clear and exhibited high contrast, while near the edge of the FOV (x = -1.8 mm and x = 1.8 mm), the reconstruction quality was degraded. After 20,000 training iterations, the cross-coherence was greatly improved both in the center and near the FOV edges.

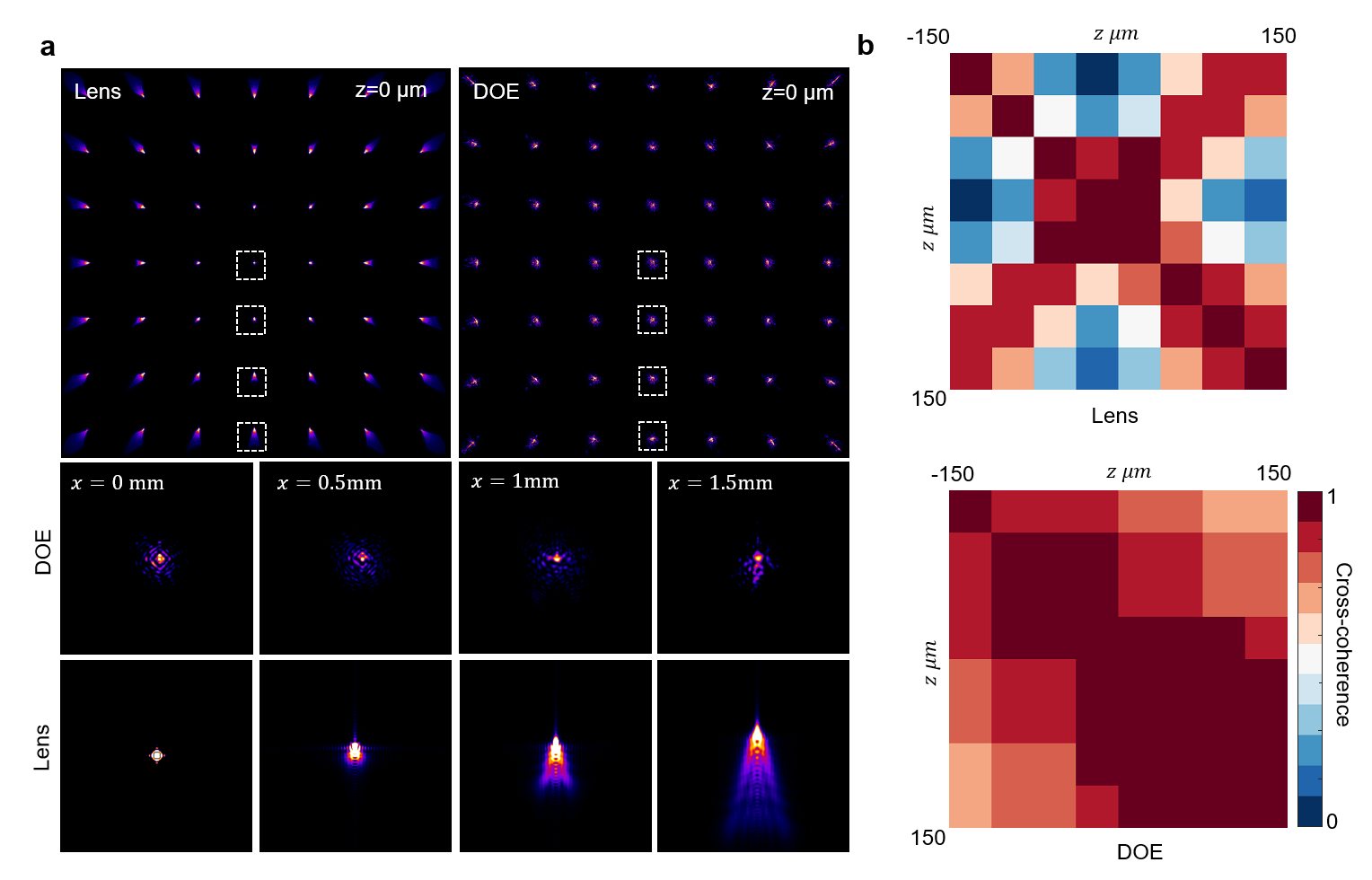

Supplementary Figure 3

Characteristics of the optimized DOE in SOMM compared to a plano-convex lens.

**a**. Top: tiled false-color plots of PSFs for different lateral positions in the focal plane for a plano-convex lens (left) and the optimized DOE (right). Bottom: Zooms into dashed boxes in (a), for different field positions.

**b**. Cross-coherence matrix between PSFs across different depths, for the plano-convex lens (top) and the optimized DOE (bottom).

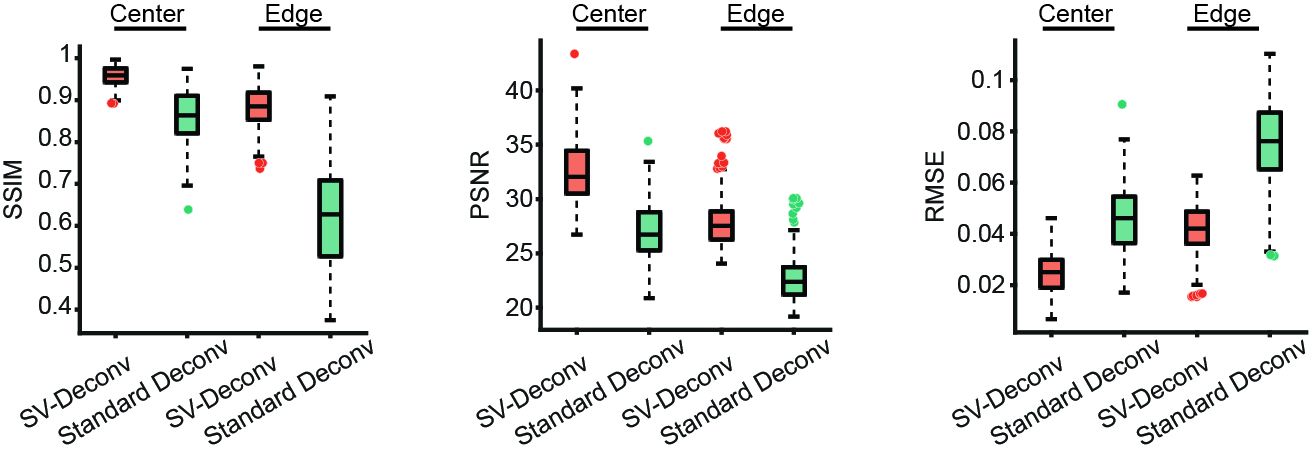

Supplementary Figure 4

Comparison of proposed SV-Deconv algorithm with standard, shift-invariant deconvolution applied to a large FOV.

Box plots of the well-known metrics structure similarity (SSIM, left), peak signal-to-noise ratio (PSNR, center), and root-mean-square deviation (RMSE, right) for the proposed shift-variant deconvolution (SV-Deconv, Methods) and a standard, shift-invariant deconvolution, each for positions in the center and near the edge of the FOV. The SV-Deconv separately deconvolves different field positions with PSFs that are specific to that position, and optimized parameters. The standard, shift-invariant deconvolution uses the central PSF to deconvolve across the entire FOV. Box centerline: median. Box shoulders: 25^th^ and 75^th^ percentile. Whiskers: Minimum and maximum excluding outliers (points). n = 159 positions near the center of the FOV and n = 197 positions near the edge of the FOV were evaluated.

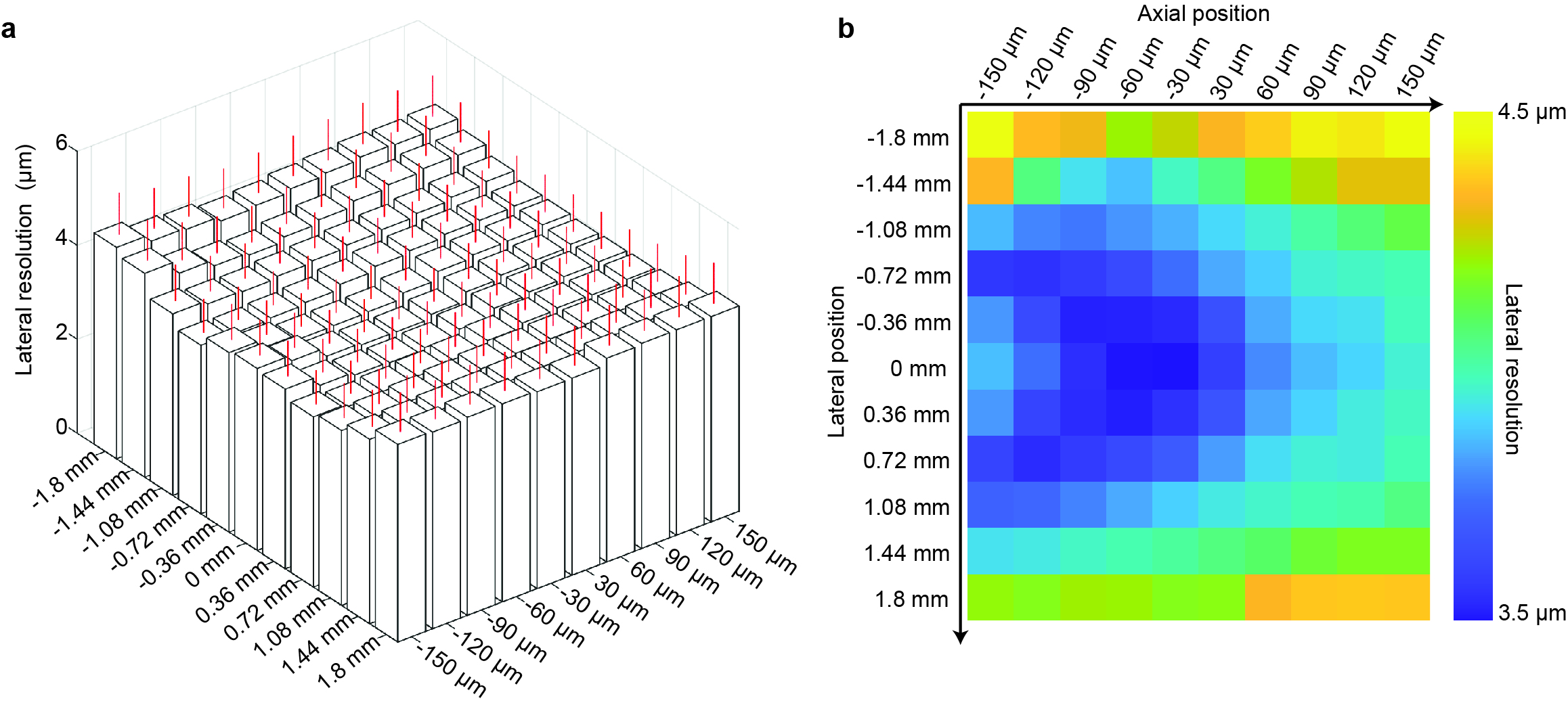

Supplementary Figure 5

Resolution of SOMM evaluated by numerical simulation.

**a**. 2D boxplot of resolution of SOMM for different lateral positions (bottom left axis, ±1.8 mm range) and different axial depths (bottom right axis, ±150 µm range). The red lines represent SD over 10 simulated, 1-µm-diameter emitters. The average resolution is 3.9 µm across the whole volume.

**b**. Heatmap of resolution of SOMM for different lateral positions and different axial positions. The resolution reaches its best value of 3.5 µm at axial position z = 0 µm and at the center of FOV, and its worst value of 4.5 µm at z = -150 and +150 µm and the edge of FOV (1.8 mm from the center).

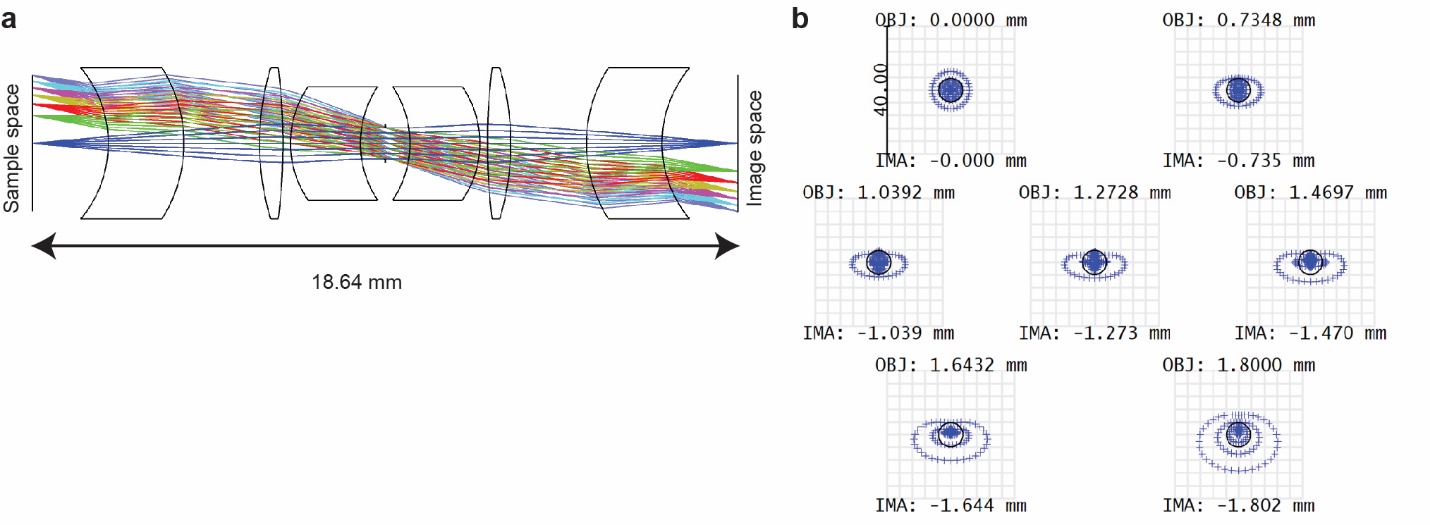

Supplementary Figure 6

Specifications of a system of spherical lenses with the same optical resolution and FOV as SOMM.

**a**. System layout and ray trace for a customized microscope with the same effective focal lens (8 mm), magnification (1×), and numerical aperture (0.1) as SOMM, but realized with spherical lenses instead of a DOE. The optical conjugation distance, i.e. distance between the sample plane and the image plane is 18.6 mm, compared to 16 mm in SOMM. The total glass volume is 88 mm^3^, which is 130 times larger than for the optimized DOE in SOMM.

**b**. Spot diagram of the spherical system at different field positions. The RMS radii of PSFs at field radius 0 mm, 0.735 mm, 1.039 mm, 1.273 mm, 1.470 mm, 1.643 mm, and 1.8 mm are 3.7 µm, 3.6 µm, 3.8 µm, 4.1 µm, 4.6 µm, 5.3 µm, and 6.4 µm, respectively.

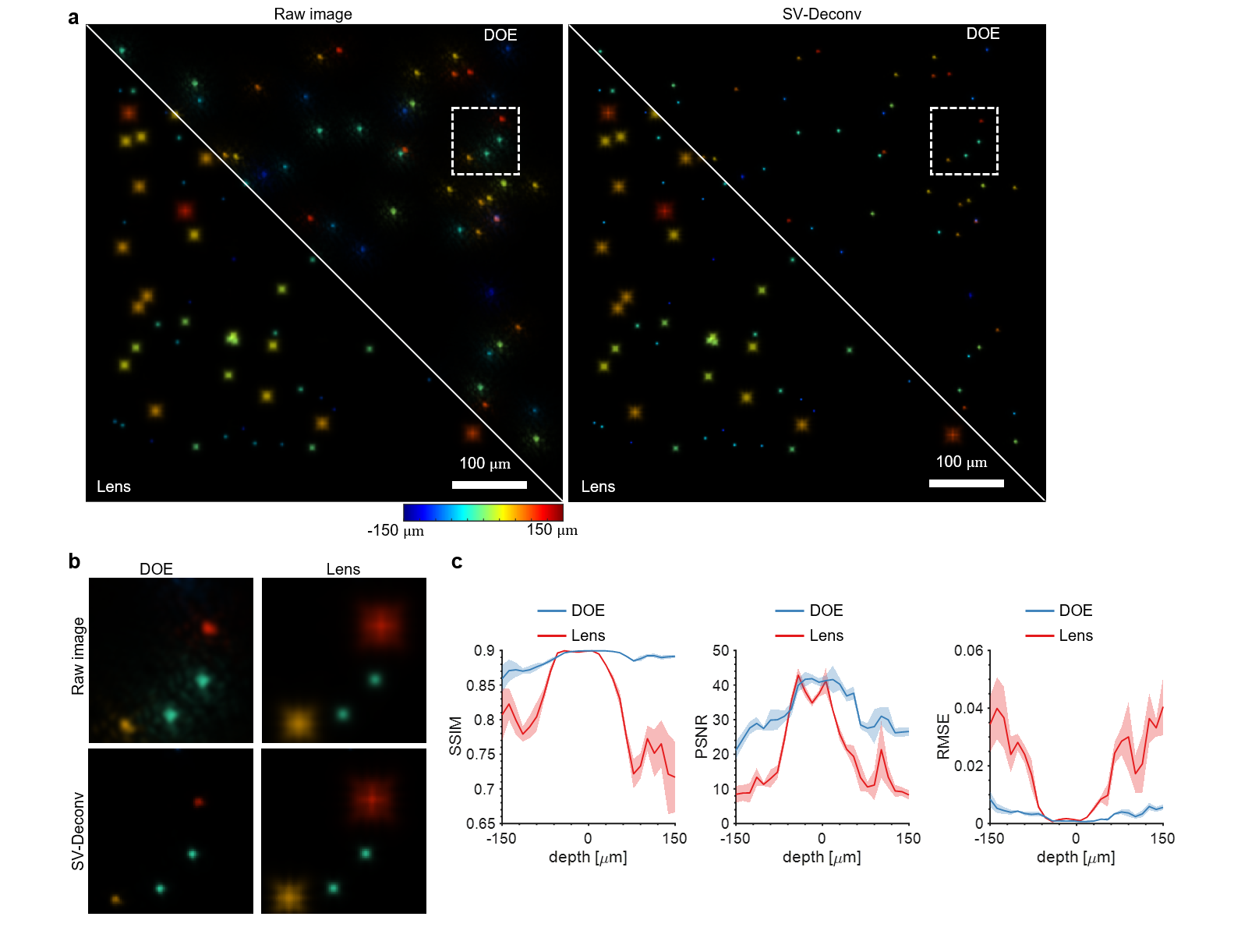

Supplementary Figure 7

Large depth of focus due to elongated PSF in SOMM.

**a**. Comparison of SOMM and a replica with a plano-convex lens in simulation, imaging emitters spread out over an axial range (defocus) of ±150 μm (Methods). Left, simulated raw images by SOMM (DOE, top right) and the replica with the plano-convex lens (Lens, bottom left) with simulated emitters randomly distributed over a 300-µm axial range. Depth of emitters indicated using false color scale. Right, SV-Deconv deconvolution results for SOMM (top right) and the lens-based SOMM replica (bottom left).

**b**. Zooms into dashed boxes in **a**. Raw (top row) and deconvolved (bottom row) images of emitters at different depths for SOMM (left column) and the replica with plano-convex lens (right column). While SOMM-imaged emitters appear similar regardless of depth, those from the lens-based replica show strong variation across depths.

**c**. SSIM, PSNR, and RMSE metrics between deconvolved images of emitters and ground truth for SOMM (DOE, blue lines) and the lens-based SOMM replica (Lens, red lines), as a function of emitter depth. SOMM achieved uniform performance across a 300-µm axial range, whereas for the lens-based replica, all metrics exhibit deteriorating performance at large defocus. Solid lines: mean. Shaded areas: SD.

Scale bars in **a**: 100 µm.

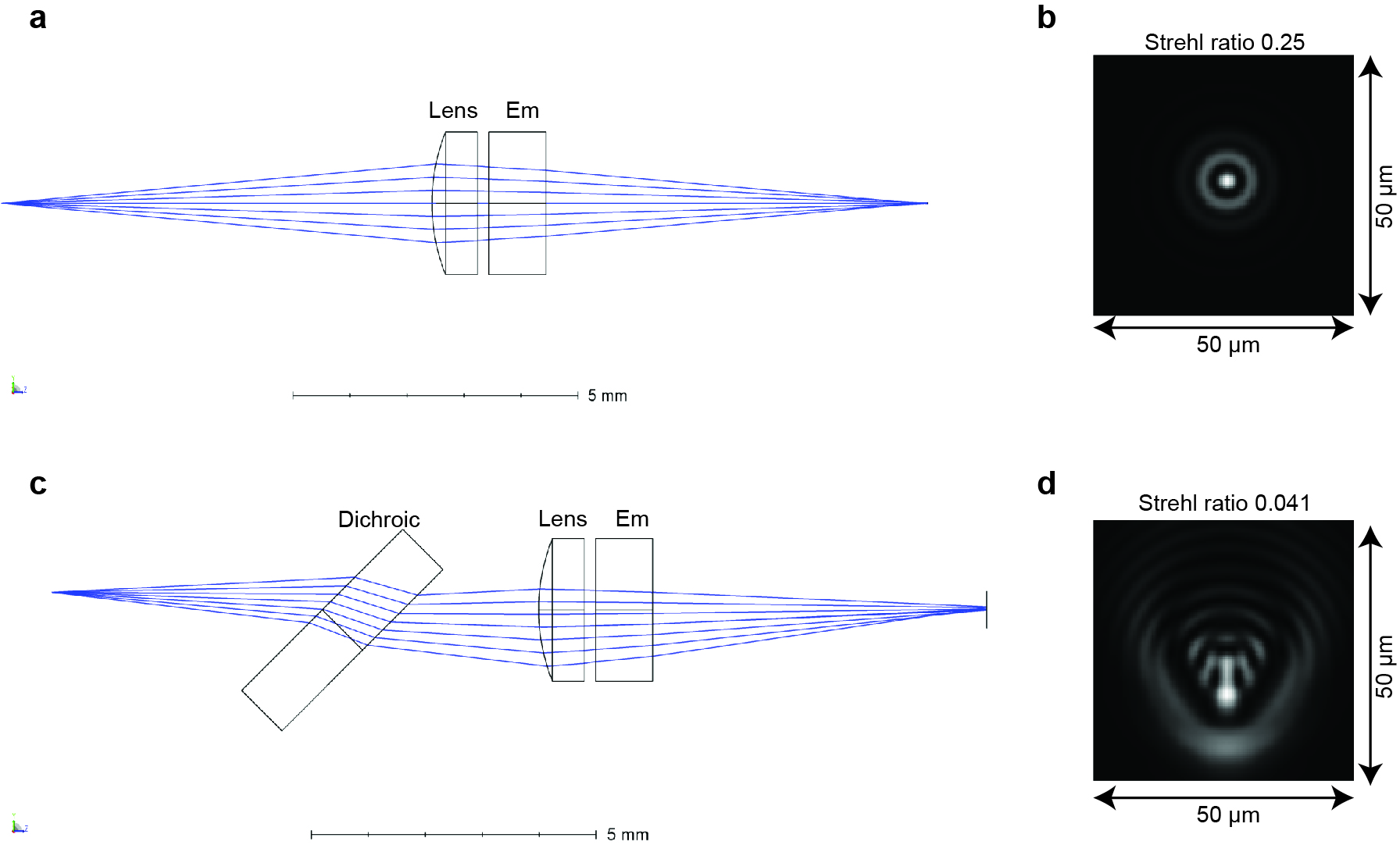

Supplementary Figure 8

Effects of imaging through a dichroic in a single-lens imaging system akin to SOMM.

**a**. A single-lens microscopic system (Edmund, #65-287), where the lens has the same aperture (1.5 mm) and similar focal length (4 mm) as the DOE in SOM, with an emission filter inserted. Such a configuration would be required for tilted illumination into the system.

**b**. Huygens PSF evaluation of the system in **a**. The Strehl ratio is 0.25.

**c**. A single-lens microscopic system (Edmund, #65-287), where the lens has the same aperture (1.5 mm) and similar focal length (4 mm) as the DOE in SOM, with a dichroic and an emission filter inserted. Such a configuration would be required for coupling illumination into the system by reflection off the dichroic.

**d**. Huygens PSF evaluation of the system in **a**. The Strehl ratio is 0.041.

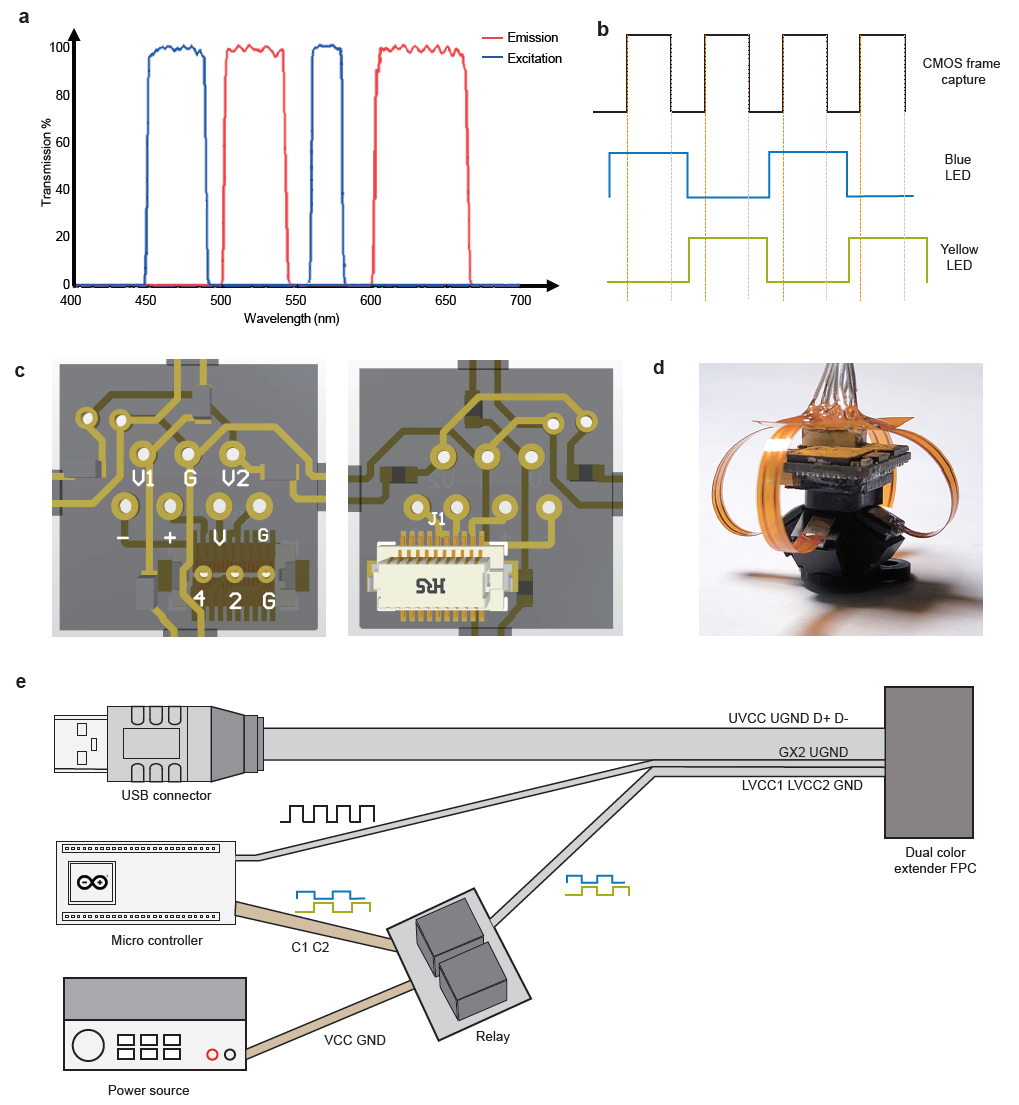

Supplementary Figure 9

Dual-color SOMM enables dual-color imaging in a compact form.

**a**. Dual-color SOMM excitation (blue) and emission (red) filter spectra.

**b**. Dual-color SOMM control signals for dual-color imaging. The control signals include the CMOS image sensor frame acquisition active signal (top), blue LED enable signal (middle), and lime LED enable signals (bottom). The specifications of LEDs can be found in Methods.

**c**. Designs of the flexible PCB for dual-color SOMM. The V1, G, V2 pins in the first row were connected to the power sources of the blue LED and the lime LED with a common ground. The -, +, V, G pins in the second row are signal and power pins of the USB2.0 connection, respectively. The 2, 4, G pins in the third row were used to receive the trigger signal for frame capture synchronization.

**d**. Photo of dual-color SOMM with flexible PCBs for connection of the LEDs.

**e**. Connection setup for dual-color SOMM. For each frame, the microcontroller sent a trigger signal to the sensor at the start of the frame (line GX2). For dual-color illumination, the microcontroller turns the blue LED on during every even frame, and the lime LED on during every odd frame. The LED control signals were sent from the microcontroller to a relay over the C1 and C2 lines, and the LEDs were powered by LVCC1, LVCC2, and GND lines. The captured frames were transmitted to the PC over an optimized thin USB cable (lines UVCC, UGND, D+, D-).

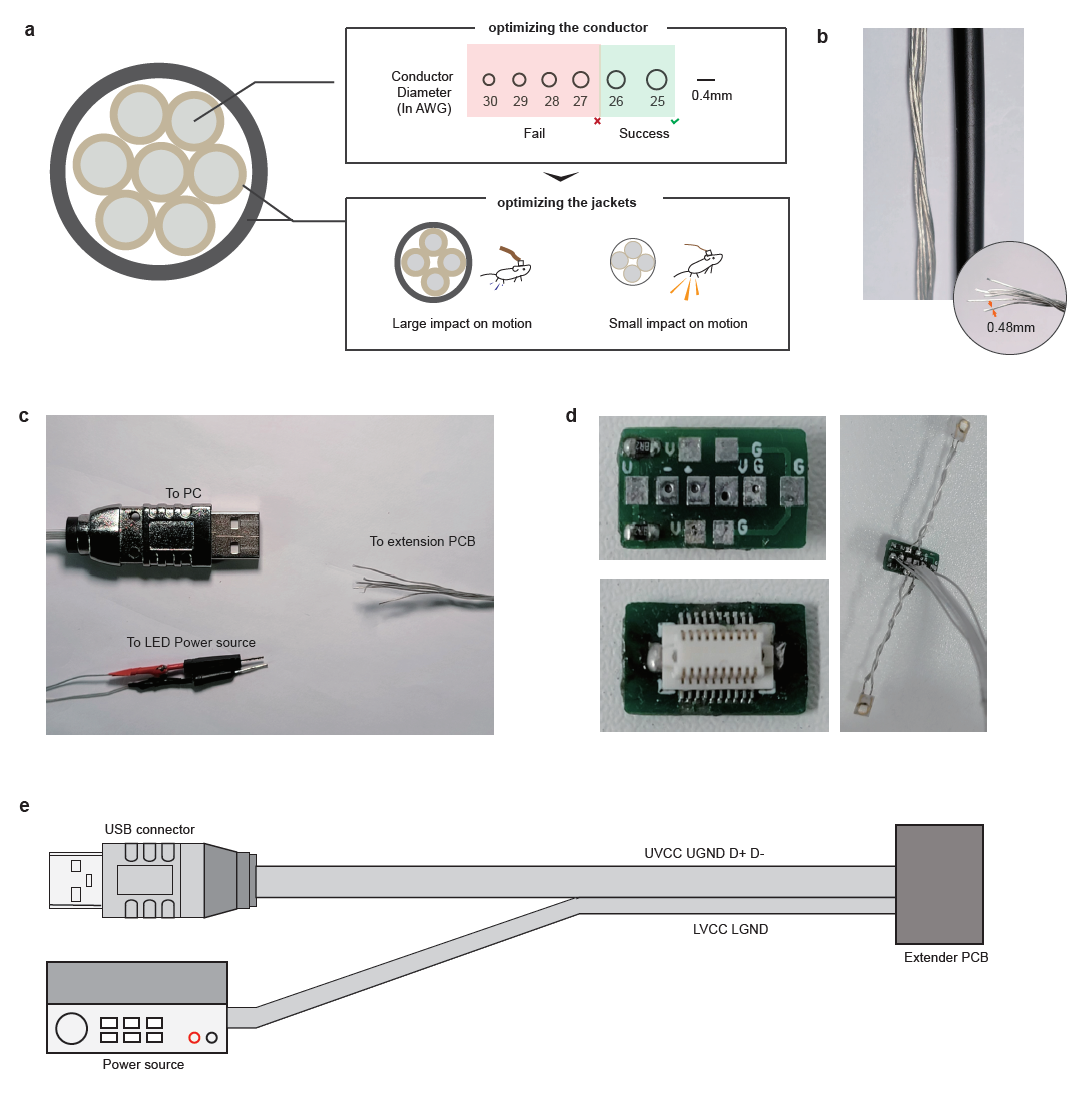

Supplementary Figure 10

Optimization of cables in SOMM.

**a**. Optimization of both conductors and jackets of the cable. First, we went through all available 6-conductor wires with the conductor diameters varying from 0.25 mm to 0.45 mm (AWG from 30 to 25, respectively), and found that only with conductor diameters larger than 0.41 mm the sensor image can be transmitted. Second, we went through all available wires with 0.41 mm conductor diameters and picked the ones with the thinnest jackets, to maximize the cable flexibility.

**b**. Photo of the final selected cable. The selected cable had 6 wires, each with an outer diameter of 0.48 mm and a conductor diameter of 0.41 mm. The overall thickness of the cable is much smaller than a standard USB cable.

**c**. Modification of a USB hub for the SOMM cable, where LED power wires and data transportation wires were integrated.

**d**. Fabricated connectors (left) and the connector with wires soldered (right).

**e**. Connection setup for SOMM. Four wires were required for transmitting signal and power for the MT9P031 sensor through the USB2.0 protocol (lines UVCC, UGND, D+, D-), and 2 wires were required to drive the illumination LED (lines LVCC, LGND).

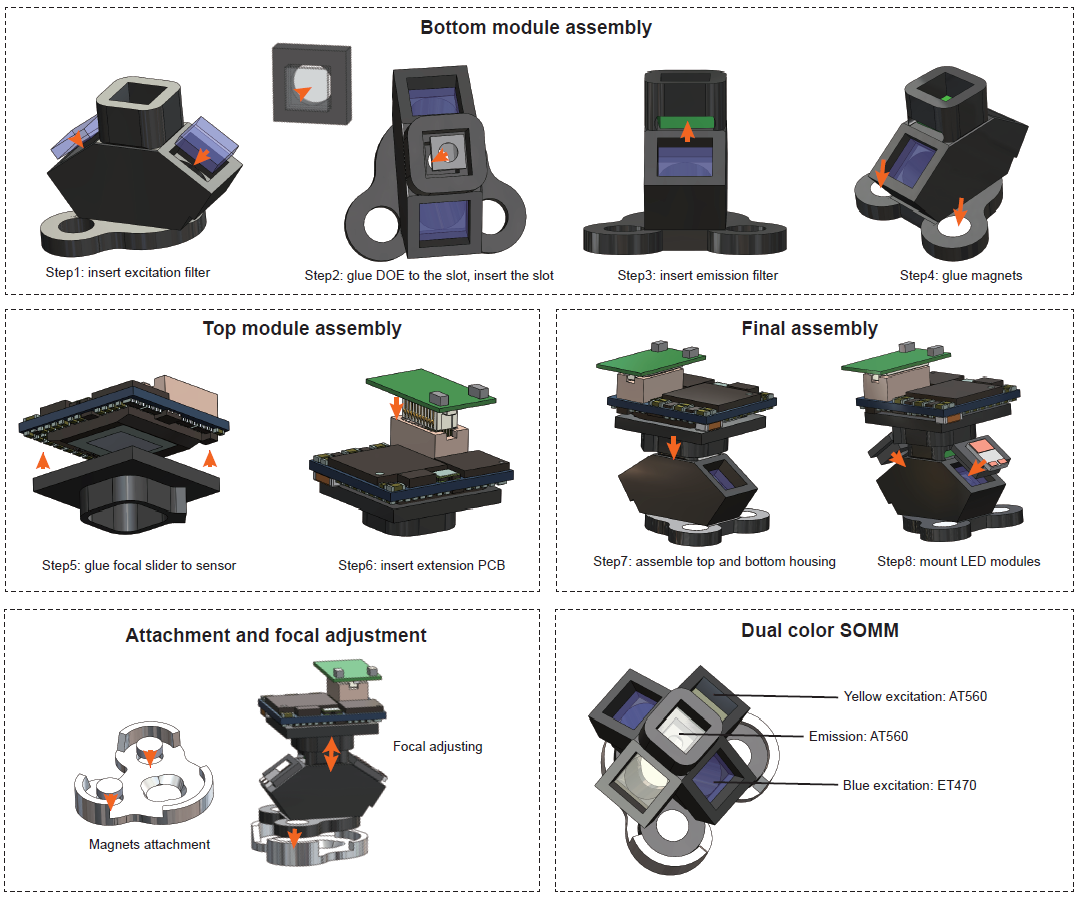

Supplementary Figure 11

Step-by-step assembly illustration for SOMM and dual-color SOMM.

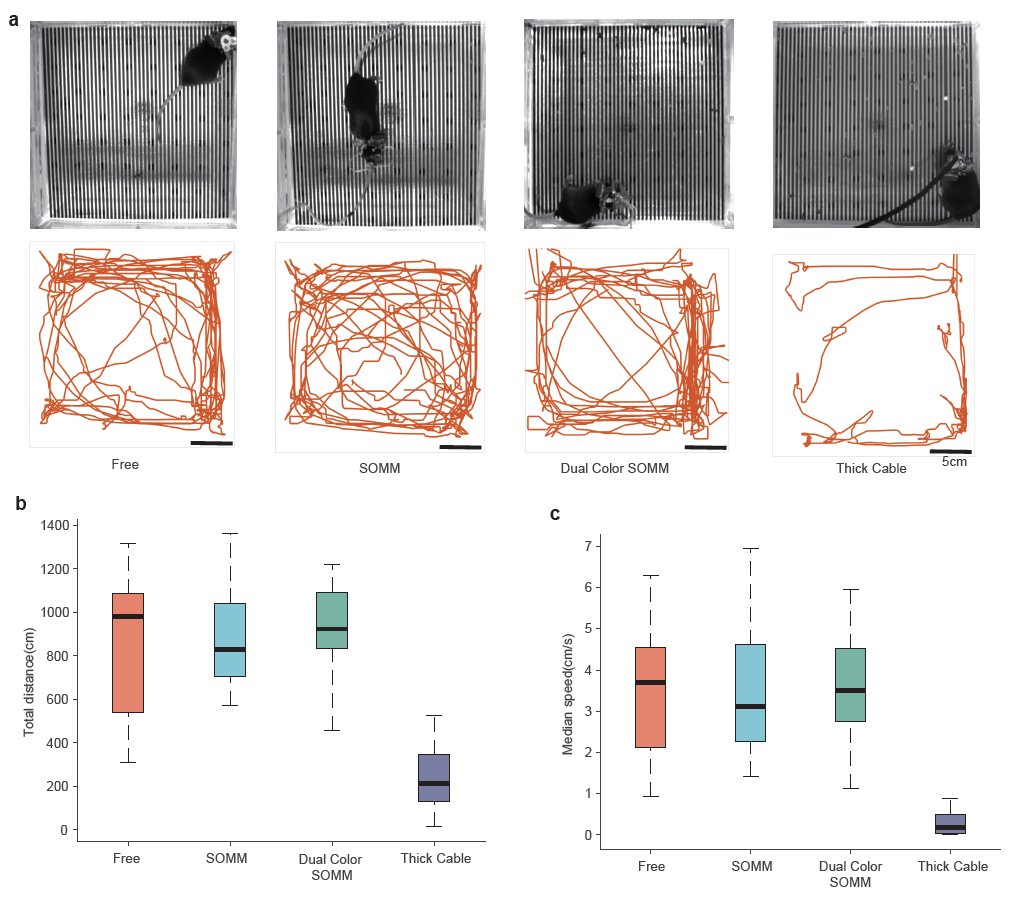

Supplementary Figure 12

Negligible effects of carrying SOMM and dual-color SOMM on mouse motion statistics.

**a**. Stills from behavior videos (top row) and extracted motion trajectories (bottom row) for mice carrying different headpieces. First column: representative mouse in homecage, with the baseplate only. Second column: mouse with baseplate and SOMM connected by the optimized cable. Third column: mouse with baseplate and dual-color SOMM connected by the optimized cable. Fourth column: mouse with aseplate and dual-color SOMM connected by the unoptimized cable (Thick cable, right). Bottom row: extracted mouse trajectories during 3-minutes of free movement.

**b**. Total distance travelled for the four conditions shown in **a**. Data from 2 trials per condition, each for three mice. Total number of trials: n = 24. Trial duration: 3 minutes. Box center line, median; Box shoulders: 75th and 25^th^ percentile; whiskers, maximum and minimum.

**c**. Median speed for four conditions listed in **a**. Symbols are the same as in **b**.

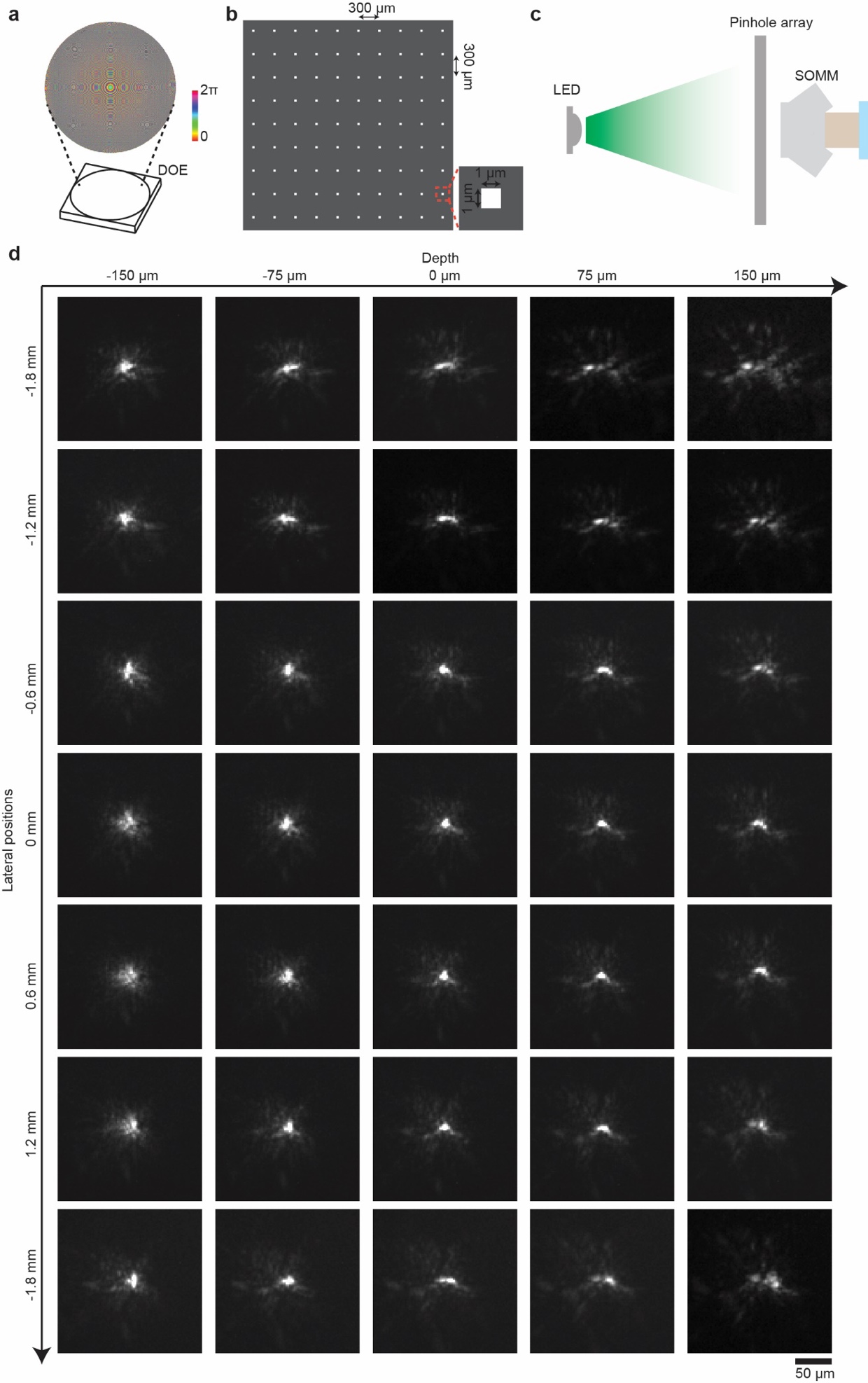

Supplementary Figure 13

PSF calibration for SOMM using a customized pinhole array.

**a**. Schematic drawing and phase pattern of the DOE of SOMM. DOE was fabricated based on the end-to-end optimized design, on a 170 µm-thickness glass substrate. The substrate was diced into a 1.5 × 1.5 mm square and installed into SOMM.

**b**. Sketch of custom pinhole array fabricated by lithography. The distance between adjacent pinholes was 300 µm, and each pinhole was 1 × 1 µm^2^. The gray areas blocked the light and the white squares were transparent.

**c**. Sketch of the PSF calibration setup. A green LED illuminated the pinhole array to generate multiple “emitters”, which were then captured by SOMM. To calibrate PSFs across an axial range, a 3-axis motorized stage was used to translate the pinhole array along the z-axis.

**d**. Experimentally calibrated PSFs of SOMM. PSFs from different lateral and axial positions are arranged in a 2D grid. The experimentally calibrated PSFs were largely similar across 3.6 mm laterally and 300 µm axially.

Scale bar: 50 µm.

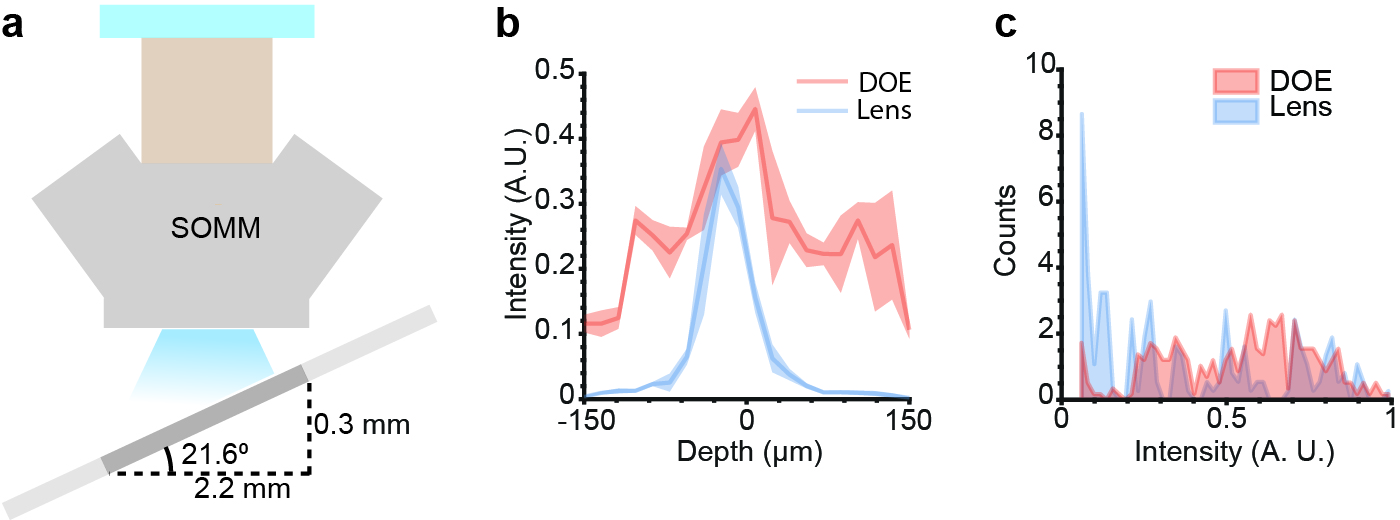

Supplementary Figure 14

Characteristics of SOMM compared to a SOMM replica with a plano-convex lens when imaging axially tilted samples.

**a**. Illustration of SOMM imaging of a tilted slide containing 6-µm-diameter fluorescent beads. The slides were tilted such that different lateral field positions were associated with different depths.

**b**. Distribution of intensity of captured fluorescent beads by SOMM (DOE, red curve) and the lens-based SOMM replica (Lens, blue curve; see Methods for the details of the SOMM replica) across a 300-µm depth range. Solid line: mean, Shaded area: SD across beads.

**c**. Histogram of intensities of captured fluorescent beads by SOMM (DOE, red curve) and the lens-based SOMM replica (Lens, blue curve) as captured across the tilted slide.

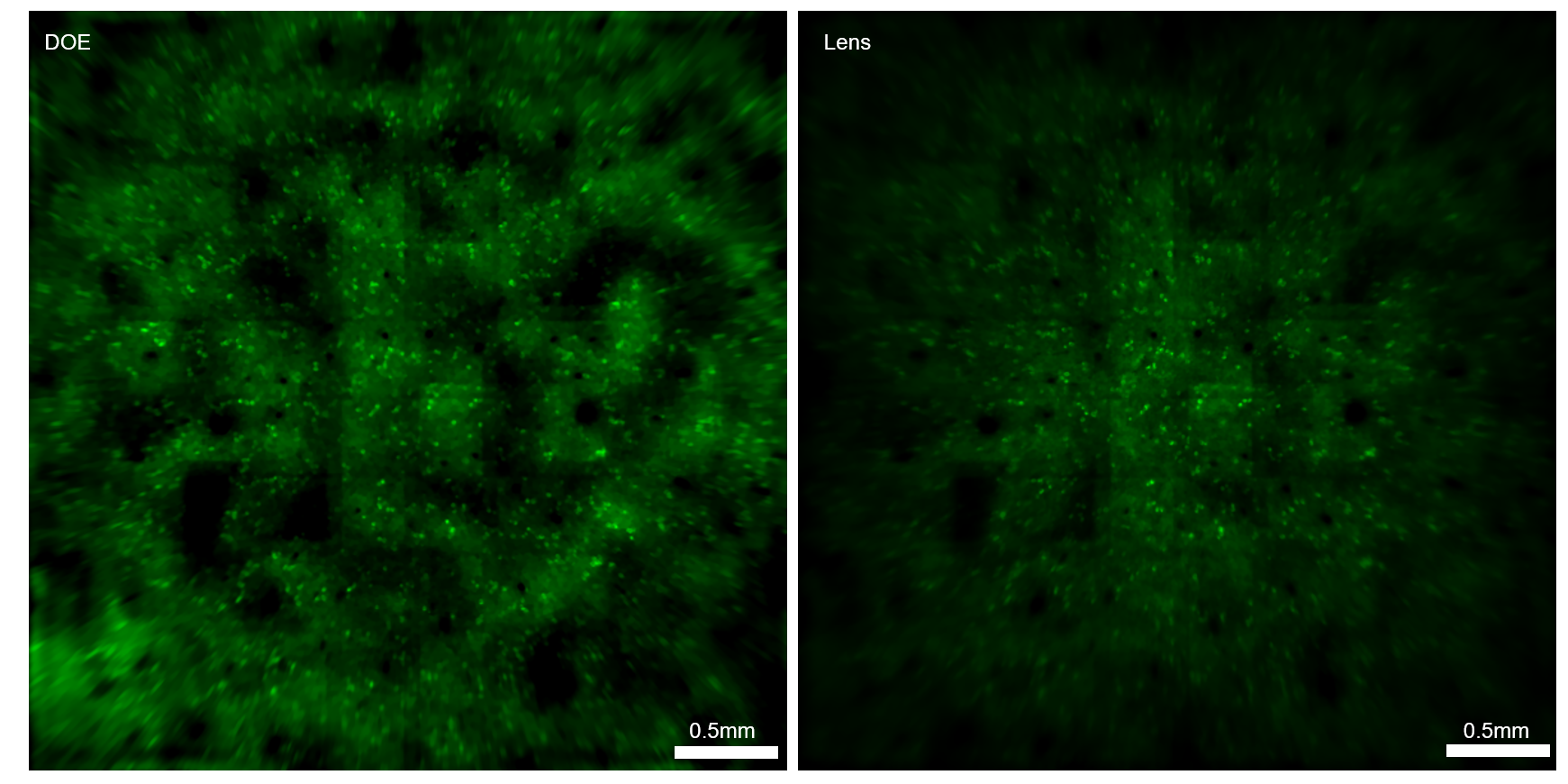

Supplementary Figure 15

Characteristics of SOMM in simulated neuronal recordings compared to a lens-based SOMM replica.

Images showing the standard deviation projection along the temporal dimension of a simulated SOMM movie capture (DOE, left) and a lens-based SOMM replica capture (Lens, right). The lens had the same focal lens and aperture size as SOMM but used a plano-convex lens rather than the optimized SOMM DOE (Methods.

Scale bars: 0.5 mm.

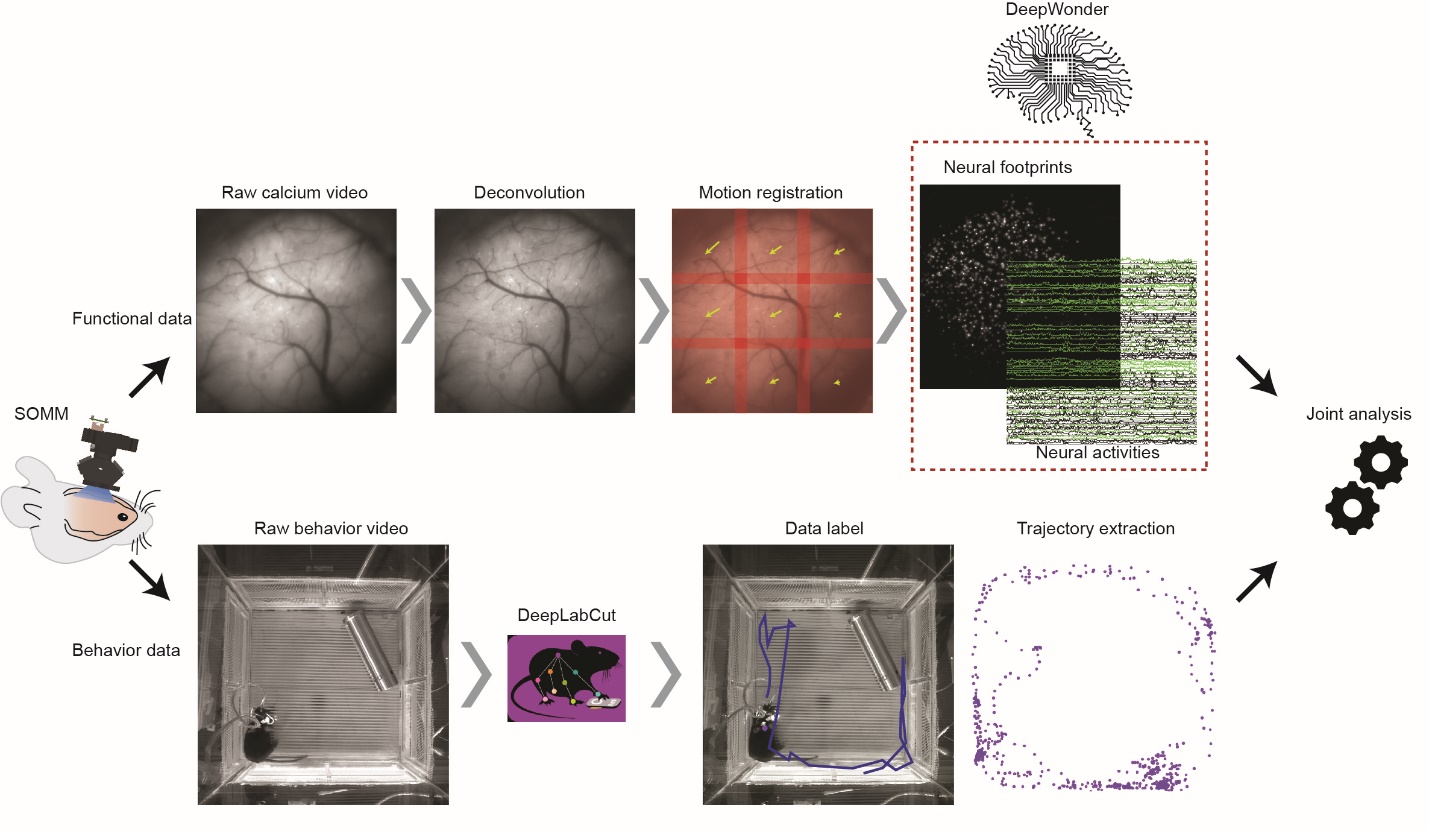

Supplementary Figure 16

Calcium imaging processing pipeline of SOMM.

The captured raw data consisted of a neuronal activity video (top) and a behavior video (bottom). The captured raw neuronal activity video was first deconvolved and motion-corrected (Methods). Then, the recently developed DeepWonder [2], which has been shown to achieve robust signal extraction across various cortical imaging methods, was applied to extract both spatial footprints and temporal signals from the video.

The captured raw behavior video was processed by the pre-trained DeepLabCut software (Methods), which resulted in motion tracking traces. Both neuronal population signals and motion traces were collected and further analyzed to characterize relations between behavior and brain activities.

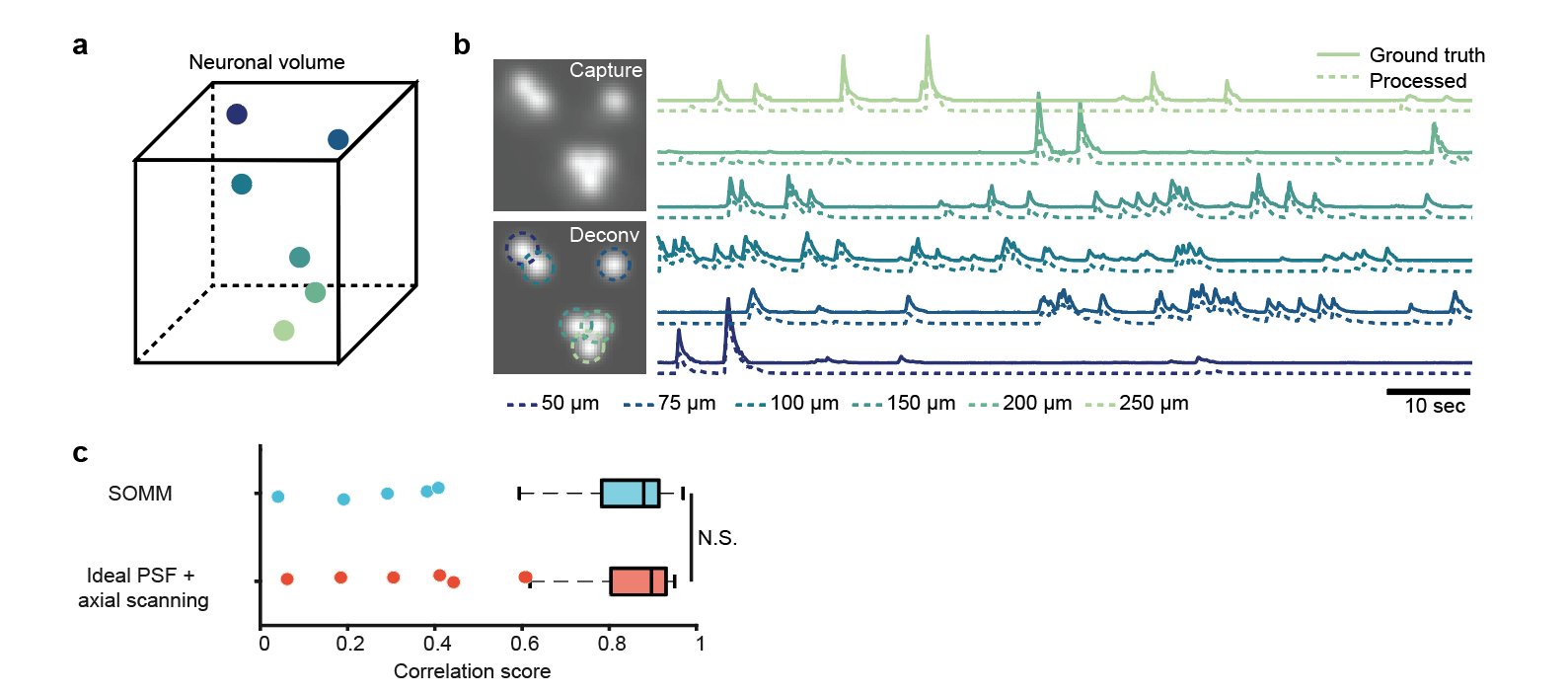

Supplementary Figure 17

Validation of calcium signal extractions in SOMM.

**a**. Illustration of a simulated tissue volume with 6 neurons at different depths (labeled by different colors) with varying lateral distances.

**b**. Left, simulated captured frame in SOMM (top) and deconvolved frame (bottom). In the deconvolved frames, neurons from different depths are marked by dashed circles with different colors. Right, ground truth signals (dashed lines) and inferred signals (solid lines), as produced by the proposed pipeline (Supplementary Figure 16), for neurons at different depths.

**c**. Temporal correlation between neuronal traces detected by SOMM (blue; 0.82 ± 0.18; mean ± SD) and an 3D scanning with an ideal, diffraction-limited PSF (red; 0.83 ± 0.17; mean ± SD). The SOMM signals were processed using the pipeline shown in Supplementary Figure 16. The signals for 3D-scanning of an ideal PSF were directly extracted from non-overlapping pixels. No significant differences were observed. N.S.: p > 0.1, two-sided Wilcoxon signed-rank test, n = 78 neurons.

Scale bar: 10 seconds.

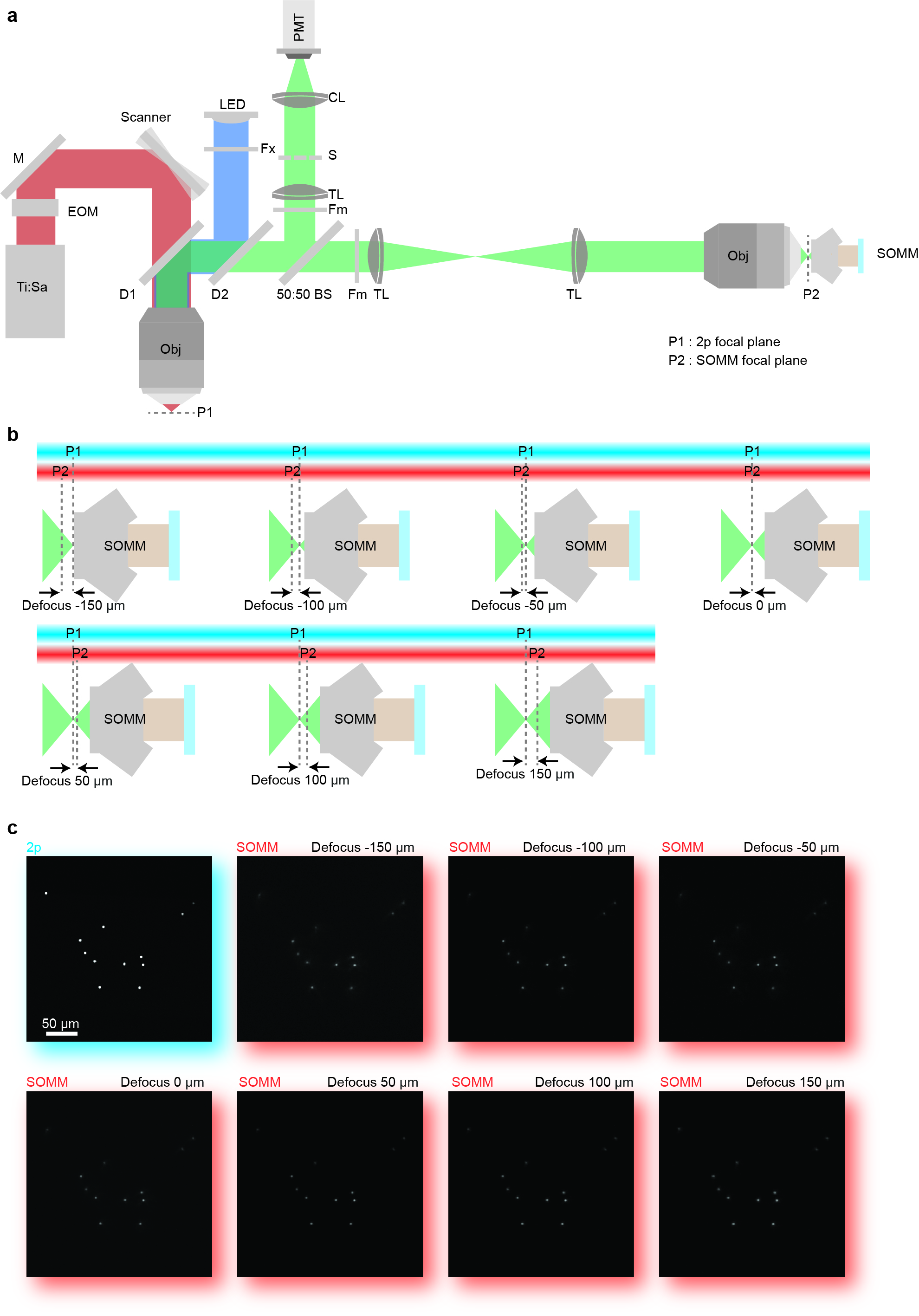

Supplementary Figure 18

System setup for temporally interleaved 2p–SOMM functional ground truth recordings.

**a**. Schematic of hybrid 2p-SOMM microscope setup used for functional ground truth recordings. LED, light-emitting diode light source; Ti:Sa, titanium: sapphire laser; EOM, electro-optical modulator; M, mirror; DM, dichroic mirror; BS, beam splitter; Fm, emission filter; Fx, excitation filter; CL, collection lens; TL, tube lens; S, triggerable shutter. SOMM was mounted after a de-magnification module that consisted of a tube lens and an objective identical to the magnification module. Under such a configuration, the 2p focal plane (P1) was optically conjugated with the SOMM focal plane (P2).

**b**. Various defocus configurations for SOMM, for verification across the DOF. We changed the position of SOMM focal plane (P2) relative to the static 2p scanning plane (P1) from -150 µm to +150 µm, which is the entire DOF of SOMM.

**c**. Representative examples of verification data captured using 2p (blue) and SOMM (red) for a range of defocus distances. We first conducted imaging of a layer of 4-µm fluorescent beads using 2p. Then, we captured the fluorescent beads for a range of defocus distances using SOMM as shown in **b,** and matched the FOV with that of 2p.

Scale bar: 50 µm.

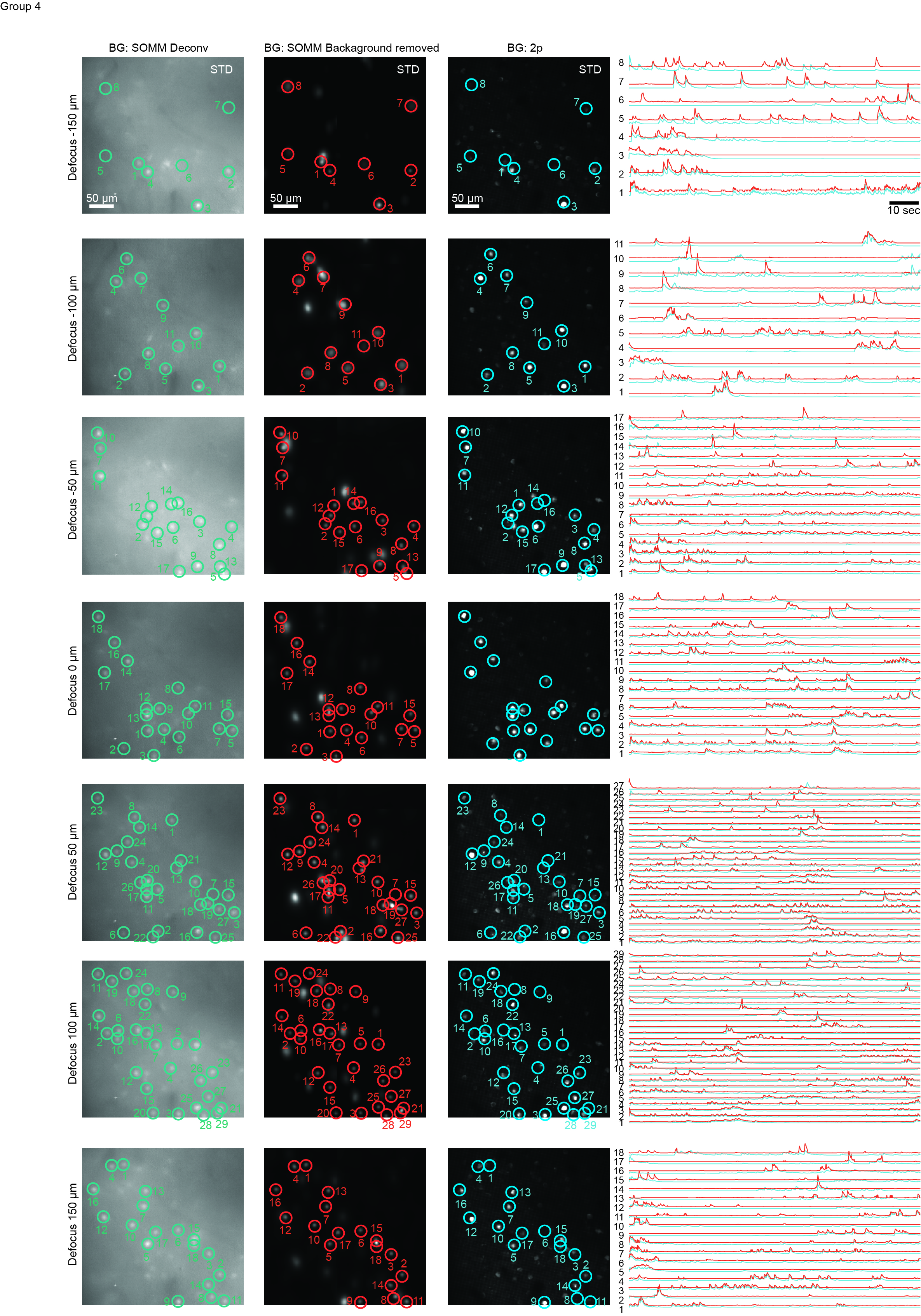

Supplementary Figure 19

Temporally interleaved 2p–SOMM functional ground truth validation.

Spatial and temporal validation of SOMM with 2p functional ground truth recordings. Temporally interleaved 2p functional ground truth was generated by interleaving SOMM frames and planar 2p microscope frames (Methods). First to third columns: standard deviation projections along time, for the deconvolved SOMM video, the background-subtracted SOMM video, and the 2p video. Rows: different defocus distances between SOMM and 2p focal planes, over a range of -150 μm to +150 μm. (Configurations illustrated in Supplementary Fig. 18). SOMM-extracted neuronal positions shown as circles overlaid on the deconvolved (first column, green) and background-subtracted videos (second column, red). Paired 2p neuronal positions were analyzed by CaImAn followed by human annotation, and were overlaid on the 2p videos (third column, blue). Fourth column: neuronal activity traces from SOMM (analyzed using the proposed pipeline in Supplementary Fig. 16; red) and 2p (analyzed by CaImAn, blue), corresponding to circles in left panels.

Scale bars: 50 µm and 10 seconds.

|  | FOV  (mm) | Resolution  (µm) | Depth of field (µm) | Frame rate (Hz) | Weight  (g) | Requires fibers? | On freely moving  mice? | Ref. |
| --- | --- | --- | --- | --- | --- | --- | --- | --- |
| Miniscope V3 | 0.7 × 0.4 | ~1.6 | ~15 | 60 | 3 | No | Yes | [3] |
| Mesoscope | 4 × 3 | ~5 | ~40 | 10 | 4.5 |  | Yes | [4] |
| FinchScope | 0.8 × 0.6 | ~1.6 | ~15 | 10 | ~4 | No | Yes | [5] |
| MiniLFOV | 3.1 × 2.3 | ~3 | ~25 | 11 | 14 | No | No | [6] |
| NINScope | 0.8 × 0.5 | ~1.6 | ~25 | 30 | 1.6 | No | Yes | [7] |
| KiloScope | 4.8 × 3.6 | ~5 | ~60 | 30 | 1.4 | Yes | Yes | [8] |
| FeatherScope | 1.0 × 1.0 | ~4 | ~40 | 30 | 1 | Yes | Yes | [8] |
| Mini-mScope | 8 × 10 | ~39 | ~150 | 30 | 3.8 | No | Yes | [9] |
| MiniLFM | 0.7 × 0.6 | 6.27 | 360 | 16 | 2.7 | No | Yes | [10] |
| CM2 | 8 × 7 | 7 | 2500 | 13 | 20 | No | No | [11] |
| CM2 V2 | 8 × 7 | 6 | 800 | 13 | >4 | No | No | [12] |
| EDoF-Miniscope | 0.6 × 0.8 | 0.9 | 70 | 30 | 2.3 | No | No | [13] |
| Bio-FlatScope | 4 × 4 | 9 | 500 | 20 | 5 | Yes | No | [14] |
| Ours (SOMM) | 3.6 × 3.6 | 4 | 300 | 16 | 2.5 |  | Yes |  |

**Supplementary Table 1: Comparison of SOMM and other miniaturized microscope.**

| **Part** | **Description / CAD file name** | **Vendor** | **Quantity** |
| --- | --- | --- | --- |
| MT9P031 | Image sensor used by SOMM | Aptina | 1 |
| CBL-MU-MH | Connection cable | Ximea | 1 |
| Extension PCB | Custom extension PCB or FPC | NA | 1 |
| DF12-20DS | Connector between sensor and extension PCB | Hirose | 1 |
| Resistor 10 | 10$\Omega$ 0603 SMD Resistors | Elecfortune | 2/4 |
| LXML-PB01-0300 | Blue excitation LED | Lumileds | 2 |
| LXML-PX02-0000 | Yellow excitation LED (dual-color SOMM) | Lumileds | 2 |
| Focus slider | Focal_Slider.SLDPRT | NA | 1 |
| Bottom housing | Mainbody.SLDPRT | NA | 1 |
| LED module | LED_holder.SLDPRT | NA | 2/4 |
| Baseplate | Baseplate.SLDPRT | NA | 1 |
| Magnet (1.5mm) | For the bottom housing, 3mm diameter, 1.5mm height | NA | 2 |
| Magnet (1mm) | For the baseplate, 3mm diameter, 1mm height | NA | 2 |
| ET470/40x | Blue excitation filter | Chroma | 2 |
| ET525/50m | Green emission filter (SOMM) | Chroma | 1 |
| ET570/20x | Yellow excitation filter (dual-color SOMM) | Chroma | 2 |
| 59022m | Dual band-pass emission filter (dual-color SOMM) | Chroma | 1 |
| DOE | Diffractive optics components | Moveon | 1 |

**Supplementary Table 2: Parts list of SOMM.**

**Supplementary Note 1: Details of end-to-end DOE optimization**

The paraxial approximation is often used to simplify the calculation of light propagation, however this however not directly suitable for mesoscale imaging. Here, to fully catch the PSF variances in large field-of-view (FOV) neuronal recording, we use the non-approximated angular spectrum method (ASM) as the propagator in the optical simulation. It is important to satisfy the sampling requirement of ASM. First, the bandwidth of the target system should satisfy $B_{total}\leq\frac{1}{2\Delta x}$ . To prevent pixel-level random phase modulation causing overly large modulation bandwith $B_{total}$ (which would prevent the applicability of ASM), we use the Zernike basis to represent the phase as $\phi=-\frac{2\pi}{\lambda}\cdot\frac{x^{2}+y^{2}}{2f}+\sum_{j} \alpha_{j}Z_{j}$ where $Z_{j}$ is the $j$-th Zernike polynomial in Noll notation and $\alpha_{j}$ is the corresponding coefficient, $f$ is the focal length of an ideal lens term that is always part of the phase profile and which we term the basic lens. The applied phase shift pattern $\phi$ has relatively small deviations from the basic lens phase, so for purposes of ASM bandwidth estimation we only calculate the bandwidth of the basic lens phase. The overall bandwidth can be calculated as [15]

$$B_{total}=\frac{L}{2 \lambda d_{1}}+\frac{L}{2 \lambda d_{2}}+\frac{\tan\left( \frac{\mathrm{FOV}}{2d_{1}} \right)}{2 \lambda}\leq\frac{1}{2 \Delta x}$$

where $\Delta x$ is the simulated step size, $d_{1}$ is the sample-DOE distance, $d_{2}$ is the DOE-sensor distance, $\lambda$ is the emission wavelength, and $L$ is the mask width. On the other hand, ASM requires a $\Delta x$ to satisfy

$$\Delta x\geq\frac{\lambda\sqrt{\left( d_{2}^{2}+\left( \frac{L_{simu}}{2} \right)^{2} \right)}}{L_{simu}}$$

where $L_{simu}$ is the simulation range, typically 2 times of the $\mathrm{FOV}$. To fully satisfy these two conditions in optimization is impractical since $L/\Delta x$ would result in memory requirements too large for a commercial GPU with 24 GB ram. For example, if $d_{1}=d_{2}=7$mm and $FOV=3$ mm, the simulation step $\Delta x$ would be as small as ~500 nm, which results in a simulation volume that is a complex array with a size of ~6000 $\times$ 6000 pixels that would need to be stored in GPU memory. Since our mesoscale optimization also has to evaluate PSFs from different sites, even a simple grid of $3\times3\times3$ positions in $x$, $y$, and $z$ would lead to simulation variables exceeding GPU memory. To implement a practically feasible optimization scheme, we exploited the fact that insufficient sampling in ASM will cause large phase inaccuracy but small intensity inaccuracy [16], which is the observable of interest in our optimizations. We therefore enlarged the sampling step $\Delta x$ such that the whole optimization model fit into GPU memory. After optimization on a GPU, we verified the performance of the optimized phase mask using a CPU and a finer grid spacing $\Delta x$ to avoid potential inaccuracy from under-sampling. Third, after the fine-grid ASM check, we discretized the optimized phase pattern using the fabrication feature size $d_{\mathrm{fab}}\times d_{\mathrm{fab}} \mu m^{2}$ and re-executed the ASM propagation to examine the effects of this discretization. In our case, the enlisted Nanoscribe Professional GT system had a frabrication resolution $d_{\mathrm{fab}}=2 \mu m$.

For DOE optimization, we used the Adam optimizer with parameters $\beta_{1}=0.5$and $\beta_{2}=0.999$. During optimization, we added random uniform noise in the range of +/- 20 nm to the DOE height map before simulating the PSF, to increase robustness to manufacturing imperfections. The training was done on a workstation equipped with an NVIDIA RTX3090 GPU with 24GB of memory.
